# Hyaluronan and Elastin-Like Protein (HELP) Gels Significantly Improve Cargo Retention in the Myocardium

**DOI:** 10.1101/2021.10.24.465557

**Authors:** Riley A. Suhar, Vanessa M. Doulames, Yueming Liu, Meghan E. Hefferon, Oscar Figueroa, Hana Buabbas, Sarah C. Heilshorn

## Abstract

Heart disease is the leading cause of death globally, and delivery of therapeutic cargo (*e.g.* cells, proteins, drugs) through direct injection into the myocardium is a promising clinical intervention. However, retention of deliverables to the contracting myocardium is low, with as much as 60 - 90% of payload being lost within 24 hours. Commercially-available injectable hydrogels, including Matrigel, have been hypothesized to increase payload retention, but have not yielded significant improvements in quantified analyses. Here, we assess a recombinant hydrogel composed of chemically modified hyaluronan and elastin-like protein (HELP) as an alternative injectable carrier to increase cargo retention. HELP is crosslinked using dynamic covalent bonds, and tuning the hyaluronan chemistry significantly alters hydrogel mechanical properties including stiffness, stress-relaxation rate, and ease of injectability through a needle or catheter. These materials can be injected even after complete crosslinking, extending the time window for surgical delivery. We show that HELP gels significantly improve *in vivo* retention of microsphere cargo compared to Matrigel, both 1 day and 7 days post-injection directly into the rat myocardium. These data suggest that HELP gels may assist with the clinical translation of therapeutic cargo designed for delivery into the contracting myocardium by preventing acute cargo loss.

## 1. Introduction

The Global Burden of Disease Study in 2017 found that 31.8% of all deaths worldwide were from cardiovascular disease, over half of which were the direct result of ischemic heart disease [1]. Since the 1960’s, the short-term (< 1 year) survival rate following a myocardial infarction (MI) has increased world-wide [2–4]. Despite these improvements, mortality from end-stage heart failure (HF) following MI is still very high, with 13% of patients being diagnosed with HF within 30 days of the initial infarction and 20-30% within the first year [5, 6]. Surgical interventions such as cardiac restraints have been shown to be insufficient at limiting HF [7, 8], and heart transplants are an unsustainable practice because of severe organ shortages [8, 9]. Direct injection of therapeutic agents into the myocardium to prevent HF can be surgically accomplished through noninvasive catheterization, making this strategy an attractive clinical option. To this end, the regenerative potential of a multitude of therapies have been studied as a means to improve cardiac function and prevent HF, including cells [10–12], gene therapy [13, 14], nanoparticles [15, 16], proteins [17, 18], drugs [19], and exosomes [20]. However, many of these therapies, which typically use saline or medium as a carrier, are severely limited because of their low retention within the contracting myocardium (**Figure 1A**) [10,12,21–24]. For example, a clinical trial studying the distribution of radio-labeled bone marrow cells injected directly into the myocardium with a liquid medium carrier found that as much as ∼90% of cells had been ejected within 1 hour [25]. Other human and animal trials studying cargo retention have shown similar results [26–29]. As cargo retention has been correlated with cardiac function and repair [30, 31], it is vital that we develop systems which improve retention post-injection.

**Figure 1.**
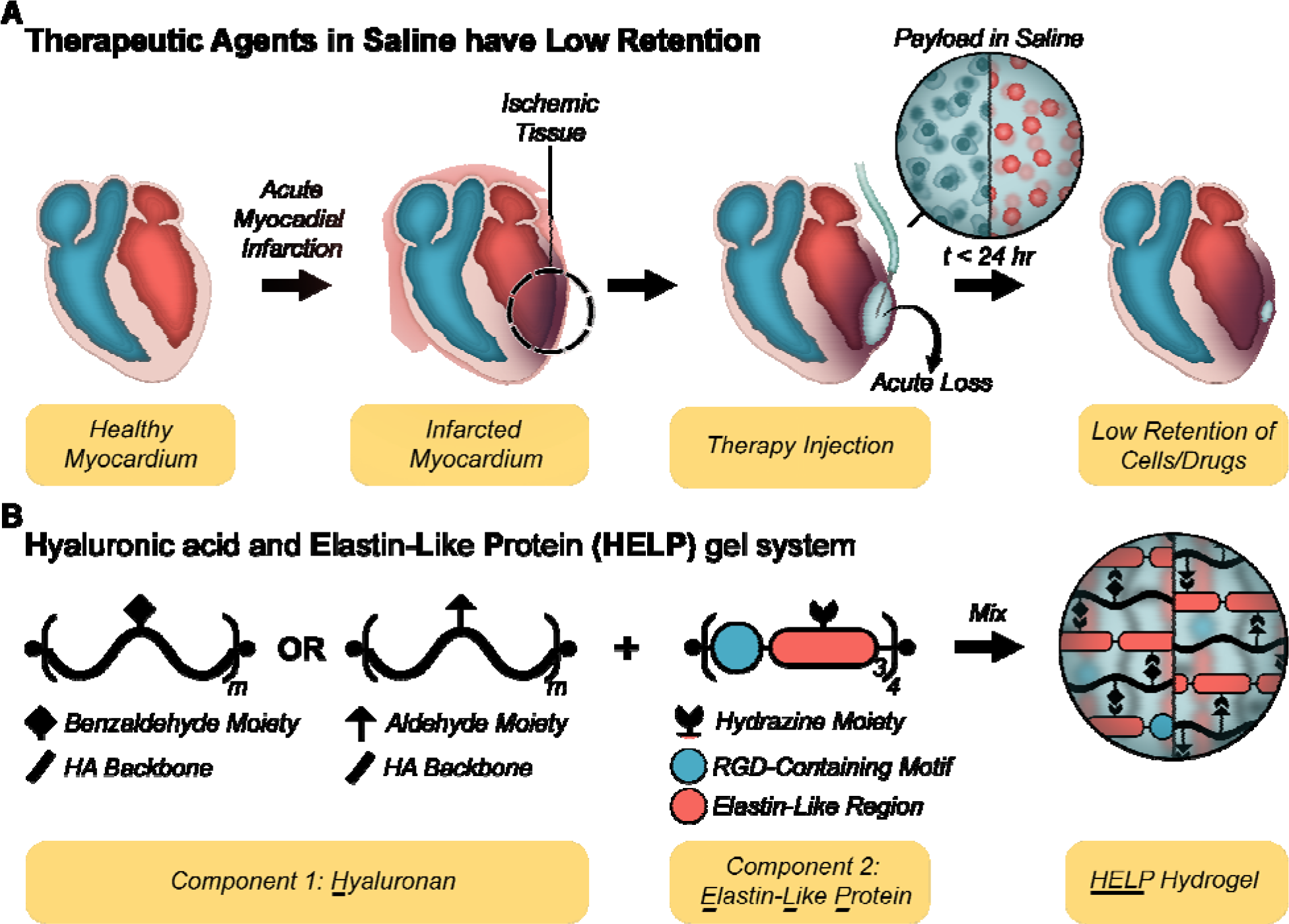
Injectable hydrogel strategy to improve myocardium retention: (A) Timeline schematic for current injectable therapies using saline. Following myocardial infarction (MI), the local tissue undergoes adverse remodeling. While several regenerative, injectable therapies (e.g. cells and/or drug-loaded particles) are being explored, they typically use saline as a delivery vehicle. Myocardial contractions cause a high degree of saline loss at the injection site, leading to low cargo retention. (B) A two-part hydrogel (termed HELP, hyaluronan and elastin-like protein) is designed to improve the retention of therapeutic agents injected into the myocardium. Hyaluronan (also called hyaluronic acid; HA) is modified to display benzaldehyde or aldehyde moieties, while the recombinant elastin-like protein (ELP) presents hydrazine moieties. These functional groups react within the syringe to form an injectable hydrogel that further stiffens in situ to localize the cargo at the injection site.

It is commonly hypothesized that injectable hydrogels may be able to increase cargo retention by physically anchoring the injected cargo within the delivery site and resisting mechanical expulsion [32, 33]. However, retention of therapeutic agents is highly variable between studies, likely due to variations in quantification methods and surgical technique [34].

Additionally, there are very few head-to-head studies comparing retention rates of different hydrogel carriers, which in light of these variable results makes direct comparisons particularly difficult. One study showed that the use of commercially available hydrogels (either Matrigel or Pluronic F127) showed qualitative reduction in acute expulsion of fluorescent microspheres compared with saline, but the differences were not statistically significant [32]. Interestingly, microsphere retention did not significantly change between the acute phase (10 minutes) and later time points (1 week), suggesting that the majority of cargo loss occurred at the time of injection and shortly thereafter. These findings correlate with clinical observations, which highlight the importance of improving retention at the time of injection [25–29]. Mechanical beating of the heart and evacuation either through ruptured blood vessels or the needle tract have been identified as key modes of acute cargo loss [29, 35]. In response, some have posited that hydrogels with more robust mechanics may better resist these mechanical challenges and therefore improve acute retention [32, 36].

Our lab has recently developed a two-part, hyaluronan and elastin-like protein (HELP) hydrogel system (**Figure 1B**) that is injectable, cell-compatible, and has independently tunable mechanical and biochemical properties [37–39]. Briefly, the hyaluronan (also termed hyaluronic acid, HA) component can be functionalized either through oxidation or conjugation chemistry to have a pendant aldehyde or benzaldehyde moiety. The elastin-like protein (ELP) is a recombinantly derived protein that contains (I) a bio-active region with a human fibronectin-derived, cell-adhesive, extended RGD sequence and (II) an elastin-like region that includes multiple lysine residues (**Figure S1)**. The lysine residues are functionalized with a hydrazine moiety, which form hydrazone bonds when mixed with aldehyde or benzaldehyde groups to spontaneously form a dynamic covalently crosslinked polymer network. In this proof-of-concept study, we expand upon this previously reported system by formulating several new HELP variants and characterizing their mechanical properties, in order to identify formulations that are ideal for use in cargo delivery to the myocardium. In particular, we demonstrate that by careful tuning of the HA component, we can significantly alter the matrix stiffness, stress-relaxation rate, and ease of injectability of HELP hydrogels through needles and catheters. We further show that HELP gels significantly increase the retention of fluorescent microspheres in the contracting myocardium *in vivo* compared to Matrigel. These data suggest that HELP gels may assist with the clinical translation of therapeutic cargo designed for delivery into the contracting myocardium by preventing acute cargo loss.

## 2. Materials and Methods

### 2.1 Expression of Elastin Like Protein

Our RGD-containing elastin-like proteins (ELP) are expressed in BL21 (DE3) pLysS *Escherichia coli* (Invitrogen, 1931008) under control of the T7 promoter, as previously described [40]. Briefly, from a frozen bacterial stock, *E. coli* is streaked across a co- ampicillin and chloramphenicol infused agar plate (25 g mL^-1^ Luria Broth (Millipore; 1.10285.5007), 15 g mL^-1^ agar (Millipore, 12177) in H_2_O; autoclaved) and incubated over-night at 37 °C. The following morning, a single colony is selected and used to inoculate a starter culture with 250 mL autoclaved growing medium (47.6 g Terrific Broth (TB; Fisher, BP9728-5)/L H_2_O) with ampicillin, and grown overnight at 37°C under constant agitation. The following morning, 20 mL of starter culture is subsequently dispersed into 12, 1L baffled flasks of growing medium and cells are grown under constant rotation at 37 °C in a shaker to an optical density (λ = 600 nm) of 0.8. λ Expression is then induced by addition of 1 mM Isopropyl β Thermo, BP1755-100) To stabilize the proteins, the orbital shakers are cooled to 32 °C. After 7 hours of expression at 32 °C, ELP-containing *E. coli* cells are collected by centrifugation, re- suspended in TEN buffer (0.1 M Tris-Cl, 0.01 M EDTA, 1 M NaCl, pH 8.0) at a final concentration of 25 g cell pellet per 100 mL, and then frozen at –80 °C overnight.

To release expressed ELP, the cell membrane is disrupted via repeated freeze-thaw cycles (3 total cycles; freezing at –80°C, thawing at 4 °C overnight). 1 mM phenylmethylsulfonyl fluoride (PMSF; ThermoScientific, 36978) and ∼ 30 mg deoxyribonuclease I (Sigma, DN25) are added after the first cycle to prevent protease activity and to degrade DNA, respectively. ELPs are further purified from the cell lysate by thermal cycling which leverages the lower critical solution temperature (LCST) behavior inherent to ELPs, and for our system is approximately ∼23 °C. After the completion of the freeze-thaw cycles, the ELP-lysate mixture is cooled to 4°C and then centrifuged at ≥ 15,000 x g at 4 °C for 1 hour. The resulting ELP-rich supernatant is collected while the ELP-lean pellet is discarded. The supernatant is then incubated at 37°C for 4 hours under constant agitation. Precipitation is further assisted by the addition 0.1 M NaCl. Combined, this induces the LCST behavior which results in a two-phase mixture of un- solubilized ELP and solution. The resulting mixture is centrifuged at ≥ 15,000 x g, 37 °C, for 1 hour and the ELP-rich pellet is then manually broken up with a small spatula to assist in re- solubilization. The mashed pellet is then mixed with ultra-pure distilled (DI) water (10 mL g^-1^) and mixed overnight on an orbital shaker at 4 °C. This thermal cycling procedure is repeated a total of 3 times. To remove exogenously added salt added during thermal cycling, the resulting solution is dialyzed against 4 L of DI water in 3.5 kDa molecular weight cutoff dialysis tubing (Spectrum Laboratories, 132592) at 4°C with constant rotation by a magnetic stir bar. The 4-L of dialysis water is routinely refreshed every 12 hours over the course of 3 days. The dialyzed product is then sterilized using a 0.22-µm syringe filter, frozen, and lyophilized for 3 days. The final lyophilized product is stored in a sterile 50-mL tube at -20 °C in the presence of desiccant to prevent water accumulation until use or down-stream modification.

### 2.2 Functionalization of Elastin-Like Protein with Hydrazine

The ELP used presently contains a central lysine (K) group within the elastin-like region. These lysine moieties were functionalized with hydrazine groups according to the following protocol, as has been reported on previously [37, 39]. First, 180 mg of ELP is dissolved in 6 mL of room-temperature anhydrous dimethyl sulfoxide (DMSO) in a 25-mL round-bottom flask. The ELP is left to dissolve completely (approximately 30 minutes to 1 hour) under constant rotation with a small magnetic stir bar. Following complete dissolution of the ELP, an additional 6 mL anhydrous N,N-dimethylformamide (DMF) is added to the reaction mixture, dropwise, using a glass serological pipette. In a separate round-bottom flask, 6 mL of anhydrous DMF is used to dissolve the following additional reagents together: (1) tri-Boc-hydrazinoacetic acid (Tri-Boc; 2 equiv:ELP amine (14 total amine/ELP); ChemImpex, 16931), (2) hexafluorophosphate azabenzotriazole tetramethyl uronium (HATU; 2 equiv:ELP amine, Sigma, 445460), and (3) 4- methylmorpholine (4.5 equiv:ELP amine; Sigma, M56557). The reagents are dissolved at room temperature (RT) by stir bar to allow for the activation of free acid groups on the Tri-Boc molecule by HATU. Next, the reagent solution of Tri-Boc, HATU, and 4-methylmorpholine is added, dropwise, to the ELP DMSO:DMF reaction-mixture which results in a final ELP concentration of 2% w/v (20 mg mL^-1^). The reaction is allowed to proceed overnight at RT under constant agitation with a magnetic stir bar.

To isolate the modified ELP, the reaction solution is added dropwise into ice-cold diethyl ether to a final volumetric ratio of 1:5 (reaction volume:ether) in a solvent-safe centrifugation tube. Precipitation of modified ELP can be further assisted by manually rocking the tube back and forth intermittently over the course of 30 minutes. ELP’s are then collected by centrifugation (>18000 x g) at 4 °C for 30 minutes in a solvent-safe centrifugation tube, decanted, and dried over-night yielding the Boc-protected ELP-hydrazine intermediate. A small sample (∼10 mg) should then be collected for downstream analysis of functionalization efficiency via NMR (below). To remove the Boc protecting groups, the ELP-hydrazine intermediate is dissolved at 2% (w/v) in a 1:1 mixture of Dichloromethane and trifluoroacetic acid with 2.5% v/v tri- isopropylsilane (Sigma, 233781) and stirred at RT for 4 hours in a vented round-bottom flask. The de-protected product (ELP-Hydrazine) is then precipitated from solution by slow addition to chilled diethyl ether using the same protocol as above. After complete separation, the solution is then centrifuged as before and air dried overnight in a well-vented fume hood. The final dried protein is then dissolved in DI water at 2% (w/v) and dialyzed against 4-L of DI water using 10 kDa MWCO dialysis tubing. The 4-L of dialysis water is routinely refreshed every 12 hours over the course of 3 days. The final dialyzed product is then sterilized using a 0.22-µm syringe filter, frozen, and finally lyophilized for 3 days in a sterile, filtered 50 mL tube (Millipore, SCGP00525).

The final lyophilized product can be stored at -20 °C in the presence of desiccant to prevent water accumulation.

NMR Quantification:

Hydrazine functionalization: dissolve ∼10 mg of the Boc-protected ELP-hydrazine intermediate in ∼750 µL deuterated DMSO and analyze by 1H NMR (500 Hz, DMSO-d6): Tetramethylsilane (TMS; 0). Efficiency of modification is determined by comparing the integrated signal of the Boc protons (δ 1.5-1.35) to the aromatic protons of tyrosine residues on ELP (δ 7.00 and 6.62, 8H each).

### 2.3 Functionalization of Hyaluronic Acid with Aldehyde and Benzaldehyde Moieties via Copper- Click Reaction

Functionalizing hyaluronan (HA) with either a benzaldehyde (HA-B) or Aldehyde (HA-A) groups via copper-click chemistry is achieved with a two-part reaction. First, commercially available 100 kDa HA (Lifecore) is functionalized via EDC chemistry to modify a specific amount of the native carboxylic acid groups to alkyne groups. Second, a commercial small molecule with either a pendant benzaldehyde group or a pendant aldehyde group is used to functionalize the alkyne groups via standard copper-click chemistry.

The procedure for alkyne modification has been described elsewhere [39]. Briefly, 200 mg of 100 kDa HA (sodium salt) was dissolved in 20 mL MES buffer (0.2 M MES hydrate, 0.15 M NaCl in DI water; pH 4.5) at a concentration of 1% (w/v). Once dissolved, sufficient propargylamine (0.8 molar equivalent; Sigma Aldrich, P50900-5G) to functionalize 12% of the available carboxylic acid groups was added directly, and immediately following 1M NaOH was used to adjust the reaction to pH 6.0. After adjusting pH, N-hydroxysuccinimide (NHS; 0.8 molar equivalent; Thermo, 24500) and 1-(3-dimethylaminopropyl)-3-ethylcarbodiimide hydrochloride (EDC; 0.8 molar equivalent; Thermo, 22980) were added sequentially and allowed to react for 4 hours at RT under constant rotation with a magnetic stir bar. The reaction was then dialyzed against 4-L of DI water using 10 kDa molecular weight cutoff dialysis tubing. The 4-L of dialysis water was routinely refreshed every 12 hours over the course of 3 days. The final dialyzed product was then sterile filtered using a 0.22-µm syringe filter, frozen, and finally lyophilized for 3 days. The final lyophilized product (HA-Alkyne) was stored in a sterile 50-mL tube at -20 °C in the presence of desiccant to prevent water accumulation. Percent modification of HA with alkyne groups can be verified by NMR (below).

Pendant aldehyde and benzaldehyde groups can be attached to HA-Alkyne via standard copper-click chemistry. Briefly, dissolve 12% modified HA-Alkyne in a sufficiently large round-bottom flask at 20 mg 0.8 mL^-1^ in 10x isotonic phosphate buffered saline (PBS; 81 mM sodium phosphate dibasic, 19 mM sodium phosphate monobasic, 60 mM sodium chloride in DI water; pH adjusted to 7.4; 0.22um filtered) with 0.85 mg mL^-1^ beta-cyclodextran. Dissolving may take up to 1 hour. After complete dissolution, degas the reaction mixture for 20-30 minutes to remove any oxygen from the reaction vial. While the HA-Alkyne is dissolving, prepare 2.4 mM copper sulfate and 45.2 mM sodium ascorbate stock solutions in DI water. After dissolving, degas both solutions for 20-30 minutes with nitrogen. Depending on the intended modification scheme, dissolve enough 4-azidobenzaldehyde (Santa Cruz Biotechnology, sc-503201) for HA-B or Ald- CH_2_-PEG_3_-Azide (BroadPharm, BP-21715) for HA-A in extra-dry DMSO (100 mg mL^-1^) for a 2- molar equivalent (reagent:alkyne). After all additional reagents have been made, sequentially add the sodium ascorbate and copper stock solutions to a final concentration of 4.52 mM and 0.24 mM, respectively. Lastly, add the –Bza or –Ald small molecule in, drop-wise. Finally, seal the round-bottom flask and degas the reaction mixture for an additional 5-10 minutes to remove any oxygen. Cover the round-bottom flask in aluminum foil and allow the reaction to proceed for 24 hours at RT under constant rotation with a magnetic stir bad.

After 24 hours, add an equal volume amount of 50 mM EDTA pH 7.0 to the reaction vial and stir for 1 hour at RT to chelate any copper in solution. Finally, dilute the solution ∼5X with DI water and sterile filter using a 0.22-µm filter to remove any precipitates. The final, dilute, solution is then dialyzed against 4-L of ultra-pure deionized water using 10 kDa MWCO dialysis tubing. The 4-L of dialysis water is routinely refreshed every 12 hours over the course of 3 days. The final dialyzed product is then sterilized using a 0.22-µm syringe filter, frozen, and finally lyophilized for 3 days in a sterile, filtered 50 mL tube. The final lyophilized product can then be stored at -20 °C in the presence of desiccant to prevent water accumulation until use

NMR Quantification:

Alkyne modification: dissolve ∼10 mg of HA-Alkyne in ∼750 µL of D_2_O and analyze by 1H NMR (500 Hz, D_2_O): TMS (0); Acetyl group (3H, δ ∼1.8); water (δ ∼4.7); alkyne (1H, δ ∼4, doublet). To determine degree of functionalization, define acetyl group as 3H for integration and compare to alkyne doublet group.

Benzaldehyde functionalization: dissolve ∼10 mg of HA-Alkyne in ∼750 µL of D_2_O and analyze by 1H NMR (500 Hz, D_2_O): TMS (0); Acetyl group (3H, δ ∼1.8); water (δ ∼4.7); triazole (1H) and benzene ring (4H) (5H total; δ ∼7.5 - 8). To quantify the degree of modification, compare the integrated peak of the triazole and benzene ring to the acetyl group.

Aldehyde functionalization: dissolve ∼10 mg of HA-Alkyne in ∼750 µL of D_2_O and analyze by 1H NMR (500 Hz, D_2_O): TMS (0); Acetyl group (3H, δ ∼i1.8); water (δ ∼4.7); aldehyde (1H; δ ∼9.8). To quantify the degree of modification, compare the integrated peak of the aldehyde group to the acetyl group.

### 2.4 Oxidation of Hyaluronic Acid

The oxidation reaction of HA has previously been reported elsewhere [41, 42]. In short,

1.5 MDa HA (sodium salt) (Millipore, 53747) is first dissolved in DI water at 4 mg mL^-1^, overnight, at 4 °C with constant agitation from a magnetic stir bar. The following day, sodium periodate (Fisher, AC198381000) is dissolved in DI water to a final concentration of 0.1 M, and subsequently added dropwise to the HA solution to a final volumetric ratio of 1:5. After addition of the sodium periodate solution, the reaction vial is covered in aluminum foil and allowed to oxidize under constant agitation with a magnetic stir bar for the desired amount of time at RT. In the present manuscript, HA was oxidized for 8 hours (HA8), 16 hours (HA16), and 24 hours (HA24). Following the desired oxidation time, ethylene glycol is added to inactivate any unreacted periodate. After one hour of deactivation, the reaction volume is transferred to 3.5 kDa MWCO and dialyzed against 4-L of ultra-pure deionized water. The 4-L of dialysis water is routinely refreshed every 12 hours over the course of 3 days. The final dialyzed product is then sterile filtered using a 0.22-µm syringe filter, frozen, and ultimately lyophilized for 3 days. The resulting products (HA8, HA16, and HA24 presently) are stored in a sterile 50-mL tube at -20 °C in the presence of desiccant to prevent water accumulation.

### 2.5 Hydrogel Formation

In the present manuscript all experiments are performed on either hyaluronan and elastin-like protein (HELP) gels with a final composition of 2% (w/v) ELP and 1% (w/v) HA, or Matrigel that is supplemented with 12.5% (v/v) PBS. All material preparation took place in a UV-sterilized fume hood and were carried out according to the following, material-specific, preparation.

For the HELP system, ELP is dissolved over night at a concentration of 4% (w/v) in sterile PBS at 4 °C under constant rotation. In tandem, HA is dissolved over-night at 4% (w/v) in 10x isotonic phosphate buffered saline at 4 °C under constant rotation. The following day, equal-volume amounts of 4% (w/v) HA and pre-chilled 1X PBS are mixed on ice by careful pipetting, ensuring no bubbles are introduced. The resulting 2% HA solution can then be mixed with an equal-volume amount of 4% ELP on ice by careful pipetting to produce a final concentration of 2% ELP and 1% HA which spontaneously crosslinks to form a gel. The relative volumes can be scaled appropriately depending on the intended application.

For the Matrigel hydrogels, commercial Matrigel (Thermo, CB-40230) is first purchased, thawed, and pre-aliquoted into appropriately sized, pre-chilled, tubes on ice. The individual aliquots are frozen and stored at –80 °C until use. The night before intended use, aliquots of Matrigel are thawed, on ice, over-night, to limit any chance for pre-gelling due to more rapid heating processes. Then, the thawed Matrigel is supplemented with 12.5% (v/v) PBS and mixed on ice by careful pipetting, ensuring no bubbles are introduced. Once warmed to RT, the resulting solution spontaneously crosslinks to form a gel. The relative volumes can additionally be scaled appropriately depending on the intended application.

### 2.6 Hydrogel Stiffness and Stress-Relaxation Measurements

Shear-rheology was carried out on a stress-controlled ARG2 rheometer (TA Instruments) using a 20-mm diameter, cone-on-plate geometry. All tests were conducted on 50- μL hydrogel samples, and heavy mineral oil was used to fill the gap between the rheometer geometry and external environment to ensure hydration over the course of all measurements. Representative values are composed of 3 – 5 replicates per gel formulation.

Samples were first crosslinked at room temperature, and a time sweep was measured over the duration at a strain of 1% and oscillatory frequency of 1 rad s^-1^ to monitor changes in mechanical properties. Gelation was first indicated by the crossover point, where the storage modulus (*G*′) was first seen to be greater than the loss modulus (*G*″). After sufficient time (HELP Gels, 30 min; Matrigel, 1 hour) all gels reached a stable ‘plateau’ modulus, and the sample temperature was then ramped to 37 °C at a ramp rate of 3 °C min^-1^. Gels were then incubated for an additional 15 min at 37 °C to ensure complete cross-linking and to allow for any additional stiffening effects from thermal aggregation of the ELP. After incubation, the shear modulus for each sample was measured at 37 °C using a non-destructive frequency sweep over the oscillatory frequency range of 0.1 - 10 rad s^-1^ at a fixed strain of 10%. Complete gelation at this stage was further confirmed by demonstration of a stable (linear) G’ over the measured frequency range, and a final stiffness value for each gel was reported as the shear moduli at 1 rad s^-1^ for consistency. Following each frequency sweep, the sample was incubated at 37 °C for 5 minutes at a low strain (1%) and frequency (1 rad s^-1^) to allow the rheometer geometry sufficient time to equilibrate. Afterwards, the sample was strained to a deformation of 10% and the stress was measured as a function of time. As an indicator of the stress-relaxation property for each gel, the time to reach half of the initial stress (t_0.5_) has been reported. To determine the t_0.5_, the measured stress values over the course of the experiment were first normalized by dividing every point over the span of the experiment by the average of the first 100 measured stress values. Then, 100 points around the halfway point (*σ*_*Norm*_ = 0.5) were selected, and a line of best fit was used to approximate t_0.5_.

### 2.7 Hydrogel Injectability Screening

Prior to *in vivo* testing, we screened our HELP gels for relative ease of injectability. To rate this injectability, we established three criterion that would need to be met in order to facilitate testing in our proposed animal experiment. These three criteria are: (1) Injectable post complete gelation (tcrosslink > 30 min), (2) easily injectable with one hand through a 30 G, hooked, needle and (3) must not have a burst injection. To assess this, ELP and HA from each of our variants: HA24, HA16, HA8, HA-A, and HA-B were prepared according to the previously stated protocols and dissolved overnight. On the day of injection 1X PBS, supplemented with food coloring (∼3 µL/mL; Market Pantry) made to match the prescribed color schemes, was used in place of standard PBS in mixing and a final volume of 50 µL of each HELP gel was loaded into the back of a 30 G syringe (Fisher, 13-689-15). Following back loading, the plunger was reinserted and the hydrogel volume was carefully moved to the base of the needle tip where it was left to crosslink at RT. After 30 minutes of crosslinking, the needle tip was carefully hooked using a pair of surgical forceps to an angle of approximately ∼90° with a ∼2 mm tip to simulate injection conditions. Each hydrogel was then injected by hand, and a video was recorded for future assessment and reference. Immediately following injection, successful completion or failure of each criterion was rated, with a check mark (?) being given for complete success and a x-mark (x) being given for a non-satisfactory result. Only HELP gel formulations that satisfied all three criteria were considered for translation to animal studies.

### 2.8 Fracture Stress Measurement

Fracture stress measurements were carried out on a stress-controlled ARG2 rheometer (TA Instruments) using a 20 mm diameter, cone-on-plate geometry. All tests were conducted on 50-μL samples prepared in accordance with the above protocols. Heavy mineral oil was used to fill the gap between the rheometer geometry and external environment to ensure hydration over the course of all measurements. Samples were first crosslinked at room temperature, and a 30- minute time-sweep was measured at a strain of 1% and oscillatory frequency of 1 rad s^-1^ to monitor changes in mechanical properties. Gelation was first indicated by the crossover point, where the storage modulus (*G*′) was first seen to be greater than the loss modulus (*G*″). Following 30 minutes of crosslinking, HELP gels were exposed to a repeated creep test wherein the % strain (γ) was measured over the course of a three-minute duration of an applied, fixed, stress (σ). Importantly, after each stress the material was allowed to relax (γ = 0) for 15 min to recover any accumulated strains. This cycle was repeated, with each subsequent stress being twice as high as the former (e.g. = 1 *p a*, 2 *p a*,4 *p a*, 8 *p a*), up until a stress that finally induced failure of the material (% strain > 2000%).

From this data, we generated plots of the viscosity as a function of the applied stress. Using Newton’s law of viscosity, we calculated a strain rate (*dγ/dt* = *γ̇*) and subsequently the apparent viscosity (*η = γ/σ*) as a function of stress (**Eq. 1**). The stress state that induced an abrupt decline in viscosity (*η* → 0*O a s*) was taken to indicate the stress at which failure occurred, and therefore the fracture stress (σ_*F*_) of the material. Reported values are composed of 4 – 5 replicates per gel formulation.

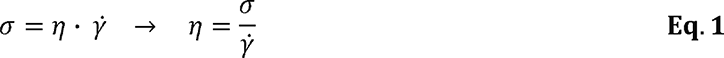

### 2.9 Post-injection Recovery and Mechanics Test

To better understand the changes in bulk-rheological properties post-injection, we performed a recovery test. Briefly, 50 µL of hydrogel were prepared as previously described and loaded into the back of a 30-G syringe. Hydrogels were allowed to crosslink for 30 minutes and then injected, by hand, onto an ARG2 rheometer (TA Instruments). Then, using a 1% strain and radial frequency of 1 rad s ^-1^, changes in storage and loss modulus were quantified using a 20 mm diameter, cone-on-plate geometry over the course of 24 hours. Heavy mineral oil was used to fill the gap between the rheometer geometry and external environment to ensure hydration over the course of all measurements. Immediately following the 24-hour recovery period, the shear modulus for each sample was measured at 23 °C using a non-destructive frequency sweep over the oscillatory frequency range of 0.1 - 10 rad s^-1^ at a fixed strain of 10%.

### 2.10 Gel Permeation Chromatography

Gel permeation chromatography (GPC) was carried out using a Dionex Ultimate 3000 instrument (including pump, autosampler, and column compartment). Detection consisted of an Optilab TrEX (Wyatt Technology Corporation) refractive index detector operating at 658 nm and a HELEOS II light scattering detector (Wyatt Technology Corporation) operating at 659 nm. The column used was a Superose 6 increase 10/300 GL. The eluent was PBS buffer with 30% (w/v) NaN_3_, 137 mM NaCl, 0.0027 mM KCl, 10 mM Phosphate pH 7.3, at 0.75 mL min^−1^ at room temperature. HA samples of 1.5MDa, 1MDa at 1 mg mL^-1^, and other analyte samples at 3 mg mL^-1^ were dissolved in ultra-pure DI water overnight and filtered through Millipore Express PES membrane filter with 0.22 pore size prior to injection. A refractive index increment (dn/dc) value of 0.165 was applied for all samples. The time frame for integration was determined by running multiple unmodified HA samples (20 kDa, 40 kDa, 60 kDa, 100 kDa, 500 kDa, 1 MDa; (Lifecore), and 1.5 MDa (Sigma)) as a reference and then applying the same integration time for our experimental samples (HA24, HA16, HA8, HA-A, and HA-B). Values represent the approximated weight average molecular weight (Mw), number average molecular weight (Mn) and the polydispersity index (PDI).

### 2.11 Dynamic Light Scattering for Measuring Hydrodynamic Radius

50-μL of each HA sample were dissolved in PBS at a final concentration of 1% (w/v).

Then, 40-μL of sample were transferred into disposable cuvettes (BrandTech, 759200) and sealed with a cap (VWR, 47744-636) to prevent any evaporation during measurements. We then performed a size measurement using the Zetasizer Nano NS, with three replicate measurements per sample. From these measurements, and assuming the viscosity of water, we were able to determine the z-average size for each sample from which we could then determine an approximate hydrodynamic radius (R_h_). The R_h_ values provided presently represent the average of three, independent, samples.

### 2.12 Labeling Elastin-Like Proteins with Cyanine-5

To aid with the visualization of our HELP hydrogels post-injection, we labeled a single lysine group on a portion of our ELPs with a cyanine-5 (Cy5) NHS molecule (excitation: 651, emission: 670; Sigma 679011) via standard NHS-ester chemistry. Briefly, ELP was dissolved in anhydrous DMSO at an initial concentration of 6% (w/v) in an appropriately sized round bottom flask at RT under constant rotation with a magnetic stir bar. Following complete dissolution of ELP, the reaction volume was diluted further with anhydrous DMF, dropwise, to a final concentration of 2% (w/v) to facilitate precipitation in diethyl ether following functionalization. Next, a 1 molar equivalent of activated Cy5-NHS ester was dissolved in anhydrous DMF (100 mg mL^-1^) and added dropwise to the 2% (w/v) ELP reaction mixture. Once added, the reaction vile was sealed with a rubber lid and purged with nitrogen gas for 5 ∼ 10 minutes to remove any excess oxygen. After purging, the reaction vile was covered with aluminum foil to prevent photo bleaching and the reaction was allowed to proceed overnight (16 hours) at RT under constant rotation with a magnetic stir bar.

To isolate the modified ELP-Cy5, the reaction volume was added, dropwise, into ice-cold diethyl ether to a final volumetric ratio 1:5 (reaction:ether) in a solvent-safe centrifugation tube. Precipitation of ELP-Cy5 was further assisted by manually rocking the tube back and forth intermittently over the course of 30 minutes which resulted in a 2-phase, ELP/supernatant, solution. ELP-Cy5 was then collected by centrifugation (>18000 x g) at 4 °C for 30 minutes in a solvent-safe centrifugation tube, decanted, and dried over-night. The dried ELP-Cy5 product is then dissolved in ultra-pure DI water at a final concentration of 2% (w/v) and dialyzed against 4- L of ultra-pure DI water using 3.5 kDa MWCO dialysis tubing. The 4-L of dialysis water was routinely refreshed every 12 hours over the course of 3 days. The final dialyzed product was then sterilized using a 0.22-µm syringe filter, frozen, and finally lyophilized for 3 days in a sterile, filtered 50-mL tube. The final lyophilized product was stored at -20 °C in the presence of desiccant to prevent water accumulation. To verify the degree of functionalization, ELP-Cy5 can be characterized via NMR (below). If the ELP molecule has been sufficiently modified (∼ 1 Cy5 molecule per ELP), ELP-Cy5 can be further modified with a hydrazine molecule (ELP-Cy5-Hyd) following the previously listed protocol and incorporated into standard HELP gels to reach a desired level of fluorescence.

NMR Quantification:

Cy5 Functionalization: dissolve ∼10 mg of ELP-Cy5 in ∼750 µL of deuterated DMSO and analyze by 1H NMR (500 Hz, DMSO-d6): Tetramethylsilane (TMS; 0). Efficiency of modification is determined by comparing the integrated signal of the lysine peak (δ 2.75-1.65; unmodified 28H) to the aromatic protons of tyrosine residues on ELP (δ 7.00 and 6.62, 8H each).

### 2.13 PEGylation of Fluorescent Microspheres

To serve as a simulated cargo, we implanted yellow-green, fluorescent, 10-µm diameter, polystyrene latex microspheres (Polysciences, 18142-2) with our HELP gels. To facilitate a monodisperse mixture and prevent clumping of our particles, we PEGylated our microspheres by functionalizing carboxylic acid groups on the microsphere’s surface with 2,000 Da Poly(ethylene)glycol (PEG) diamine molecules (Sigma, 753084) via EDC chemistry. This protocol has been reported previously [43]. First, the microsphere stock solution was vigorously vortexed for 60 seconds to ensure a homogenous mixture. After vortexing, 100 µL of 2.6% (w/v) carboxylated microspheres stock suspension was transferred to a 1.5 mL tube and subsequently centrifuged at 9,000 x g for 3 minutes to pellet the suspended microspheres. The storage buffer was carefully decanted, and then the pelleted microspheres were resuspended with 200 µL of 50 mM MES buffer, pH 6.0, and gently washed with careful pipetting. Washing is repeated for a total of three cycles, and the microspheres are finally re-suspended in 200 µL MES buffer. Next prepare: (1) a 2 mM solution of bifunctional, amine-terminated PEG in 100 mM bicarbonate buffer, (2) 200 mM EDC (Thermo, 22980) suspended in 50 mM MES buffer, and (3) 500 mM sulfo-NHS (Thermo, 24510) in 50 mM MES buffer. To the microparticle suspension, sequentially add 2 µL NHS (final: 5 mM) and 2 µL EDC (final: 4 mM) and vortex vigorously to mix. Allow the solution to incubate for 30 minutes at RT on a constant rotator to allow for activation of the carboxylic acid groups while preventing settling of the microspheres. Following 30 minutes of activation, add 200 µL PEG-diamine (final: 50mM) and mix well by vortexing. Allow the solution to incubate for 30 minutes on a constant rotator at RT to allow for PEGylation. Then, quench the reaction by addition of 400 µL 200 mM glycine in PBS (final: 100 mM) and mix well by vortexing. Incubate for an additional 30 minutes at RT on a constant rotator to ensure the reaction has completely stopped.

To purify the PEGylated microspheres, centrifuge the solution at 9,000 x g for 3 minutes and carefully decant the supernatant by pipetting. Then, re-suspend the microspheres in 800 µL ultra-pure DI water and mix by gentle pipetting. Repeat this process of centrifugation and re- suspension, each time lowering the suspension volume by 200 µL. After the 4^th^ wash (re- suspending in 200 µL ultra-pure DI water), perform two additional washes, re-suspending the microspheres in 200 µL ultra-pure DI water. Finally, centrifuge the microspheres once more at 9,000 x g for 3 minutes, carefully aspirate the supernatant, and re-suspend in ultra-pure DI water to a desired concentration. Presently, our microspheres were suspended in 130 µL DI water to achieve a final concentration of 2% (w/v). After PEGylation, microspheres can be stored in suspension at 4 °C, in the dark to avoid photobleaching, until the intended application.

### 2.14 Preparing Implantation Stock Materials

For our retention study, all hydrogels (HA16, HA24, Matrigel) were injected with a 1.25 mg mL^-1^ final concentration of 10-µm, PEGylated, fluorescent microspheres prepared according to the previously stated protocol. All HELP gel injections were delivered as 50 µL volumes with a final concentration of 2% (w/v) ELP-hydrazine (20% (v/v) ELP-Cy5-Hydrazine) and 1% (w/v) HA. Matrigel injections were delivered as 50 µL volumes with a final concentration of 87.5% (v/v). To limit the possibility of batch-to-batch variability as a side-effect of material preparation, all materials used for implantation were pre-aliquoted into injection-size volumes, stored frozen at –80 °C, and thawed overnight on ice the night before intended use. Gel stocks that were not used after thawing were discarded to reduce any variations introduced by freeze-thawing. Additionally, the mixing procedure for material and microspheres was made uniform for each system to further limit variability (below).

HELP Gels:

HA (HA16, HA24) was dissolved overnight in sterile 10X PBS to a final concentration of 4% (w/v). The following day, HA’s were aliquoted into 18-µL volumes in individual, sterile, 250-µL plastic tubes and stored frozen at –80 °C. ELP-Hydrazine and ELP-Cy5-Hydrazine were additionally dissolved overnight in sterile 1X PBS to a final concentration of 4% (w/v). The following day, ELP-Hydrazine and ELP-Cy5-Hydrazine were blended together 4:1 (ELP- Hydrazine:ELP-Cy5-Hydrazine; final: 20% labeled), and the resulting solution was aliquoted into 30-µL volumes in individual, sterile, 250-µL plastic tubes and stored frozen at –80 °C. Finally, for each injection set, a 15-µL volume of sterile 1X PBS was additionally aliquoted into individual, sterile, 250 µL plastic tubes and stored frozen at –80 °C. The night before surgery, a single aliquot of (1) 20% Labeled ELP-Hydrazine, (2) HA, and (3) PBS, per injection, was removed from storage and left to thaw, on ice, overnight. The day of surgery, injection volumes for HELP gels were prepared according to the following. First, 5 µL of 20 mg mL^-1^ stock PEGylated fluorescent microspheres was mixed by careful pipetting with 15 µL thawed, sterile, PBS (final: 5 mg mL^-1^ microspheres). Next, 18 µL of the microsphere solution was mixed by careful pipetting with 18 µL thawed, sterile, HA (final: 2% (w/v) HA; 2.5 mg mL^-1^ microsphere). Finally, 30 µL HA- microsphere solution was mixed with 30 µL of thawed, sterile ELP and by careful pipetting (final: 2% (w/v) ELP, 1% (w/v) HA, 1.25 mg mL^-1^ microspheres) and 50 µL of the HELP-microsphere solution was then back-loaded into a 30 G syringe and allowed to crosslink for > 30 min at RT prior to careful injection into the myocardium.

Matrigel:

Fresh Matrigel was thawed, and pre-aliquoted into 49-µL volumes in individual, sterile, 250-µL plastic tubes and stored frozen at –80 °C. Additionally, for each injection set, 5 µL of sterile PBS was aliquoted into individual, sterile, 250-µL plastic tubes and stored frozen at –80°C. The night before surgery, a single aliquot of (1) Matrigel and (2) PBS, per injection, was removed from storage and left to thaw, on ice, overnight. The day of surgery, volumes for Matrigel injections were prepared according to the following. First, 5 µL of 20 mg mL^-1^ stock PEGylated fluorescent microspheres was mixed by careful pipetting with 5 µL thawed, sterile, 1X PBS (final: 10 mg mL^-1^ microspheres). Then, 7 µL of the microsphere solution was mixed by careful pipetting with 49 µL thawed, sterile, Matrigel (final: 87.5% (w/v) Matrigel, 1.25 mg mL^-1^ microspheres) and 50 µL of the Matrigel-microsphere solution was back-loaded into a 30 G syringe and allowed to crosslink for > 1 hour at RT prior to careful injection into the myocardium.

### 2.15 Surgical Procedure, Animal Post-Op Care, and Euthanasia

This protocol was approved by the Stanford Administrative Panel on Laboratory Animal Care. The National Institutes of Health (NIH) guidelines for the care and use of laboratory animals were observed (NIH publication no. 85-23 Rev. 1985). All animals were purchased from Charles River and caged in gender-matched pairs for the duration of this protocol. Food and water were administered ad libitum, and their environment was temperature and humidity controlled with a 12-hour on, 12-hour off light schedule. Animals were checked every day over the course of the experiment for any signs of stress or injury to ensure a safe environment. Animals that showed evidence of distress or injury were euthanized.

First, mixed-gender Sprague Dawley rats (10-week-old, n=38; 19 19♀) were anesthetized using 3% isoflurane mixed with 1% oxygen for 10 minutes, and then manually intubated using a silicone, 16-G catheter sheath (Santa Cruz Biotech, sc-360107). Rats were then hooked up to a mechanical respirator (Harvard Apparatus, 55-7040) with 1 – 2% isoflurane and 1% oxygen to allow for mechanical ventilation (∼2 - 3 mL push volume, 80 BPM). Once steady mechanical ventilation had been established, a toe-pinch was used to verify complete anaesthetization. All procedures were carried out on a sterilized, metal board warmed with a heating pad (42 °C) for the duration of the procedure. The left portion of the chest was cleared of all fur and a left thoracotomy was performed to expose the heart. Once the wall of the left ventricle was identified by the presence of the descending coronary artery, an injection of 50 µL of hydrogel (prepared in accordance with the previously listed procedure) was administered. All injections were carried out by hand, and the needle tip was left inside the heart wall for approximately ∼10 seconds following injection to minimize the degree of reflux (expulsion of the material from the needle tract). All injections were recorded with the date and animal identifier for later review and scoring. After stable cardiac rhythm had been recovered, the chest cavity was closed with 4-0 absorbable sutures (Thermo, NC1656516) and the negative pressure of the chest cavity was restored by manually evacuating the heart of air with a 10-mL syringe affixed with a silicone catheter sheath. Finally, the chest muscles and skin were closed using 4-0 absorbable sutures and the skin was cleared of any blood or debris with sterile saline.

Following skin closure, rats received subcutaneous injections of buprenorphine SR (1.5 mg kg^-1^) as a long-acting partial opioid agonist pain reliever, bupivacaine (0.25%) as a local anesthetic, and carprofen (0.5 mg mL^-1^) as a non-steroidal anti-inflammatory. The isoflurane gas was then cut off and the animal was mechanically ventilated with 1% oxygen until they woke and removed themselves from the intubation tube. Animals were then placed into a warm recovery chamber with circulating 1% oxygen for an additional 15 – 30 minutes and monitored for signs of movement and recovery, after which the rat was returned to their housing which was warmed with a heating pad.

Rats were inspected daily up until euthanasia and monitored for general wellness, post- operative complications, infections of the skin and superficial muscle layers, and wound dehiscence. For 3 days post-surgery, rats received a daily injection of carprofen (0.5 mg mL^-1^) while under light passive inhalation anesthesia (1.5% isoflurane). If any animal was found in catatonia, immobile/paralyzed, and/or lost more than 10% of its pre-operative body weight, it was euthanized as per the Stanford Administrative Panel on Laboratory Animal Care.

Prior to euthanasia, rats underwent inhalation anesthesia using 3% isoflurane within a knock-down chamber and were then transferred to a standard nose cone. Their tails were cleaned using sterile surgical gauze and 70% ethanol. The lateral veins were then visualized by placing the tails in a beaker of warm water to promote vasodilation. Once the veins were adequately visualized, rats were euthanized via intravenous injection of potassium chloride (90 mg mL^-1^) using a 26-G needle. If visualization of the lateral tail veins was unsuccessful, rats were euthanized via an interperitoneal injection of Beuthasol, a commercial pentobarbital sodium and phenytoin sodium solution (150 mg kg^-1^) using a 23-G needle.

### 2.16 Heart Dissection and Fixation

Following euthanasia, the chest was shaved of all fur and cleaned using 70% ethanol. The skin, underlying muscle, and bones of the rib cage were removed, and the exposed heart was removed using surgical scissors. The outside of the heart was washed with fresh, sterile, 1X PBS, and then manually perfused by flushing the left and right ventricles with 1X PBS and subsequently manually evacuating them with gentle prodding. This process of manual perfusion was carried out until no evidence of blood was visible in the evacuated PBS. After perfusion, a 5-mm section of the heart, roughly centered around the injection site, was removed using a pair of fresh razor blades and a heart matrix sectioning tool (Zivic Instruments, HSRS001-1). After sectioning, the 5-mm section of heart was immersed in 40 mL of freshly prepared 4% paraformaldehyde and fixed for 72 hours at 4 °C. After fixation, the tissue section was transferred to fresh, 30% sucrose in PBS for 72 hours. Samples were stored in 30% sucrose until embedding and sectioning.

### 2.17 Tissue Embedding, Gelatin-Coating Slides, and Tissue Sectioning

All tissue sections were embedded in optimal cutting temperature (O.C.T.; Thermo, 23- 730-571) by snap freezing. First, a metal bowl was filled with 100% ethanol and rapidly cooled by chilling with dry ice. Meanwhile, tissue sections were removed from 30% sucrose storage containers, dried carefully, and then immersed in O.C.T. media in a disposable embedding mold (Fisher, 1380). Any air pockets trapped within the ventricles were carefully evacuated using a pipette, and the tissue section was centered with a consistent top-down orientation. Molds with tissue and O.C.T. media were then carefully placed into chilled, 100% ethanol, such that contact with the walls of the container was maximized but no ethanol spilled over into the tissue mold. Then, single pieces of dry ice were placed over the top of the mold both to hold the mold in place and to further assist in freezing. Each sample was allowed to snap freeze for ∼15 minutes to ensure complete freezing. Once the sample block was frozen (indicated by a transition to opaque white) samples were then stored in a –80 °C freezer until sectioning, but at a minimum of 24 hours.

To maximize tissue adhesion to our slides, we gelatin coated microscope slides (Thermo, 643203) using commonly reported methods. In short, microscope slides were first cleaned with 70% ethanol. Then, in batches, slides were immersed into gelatin-coating media (2% (w/v) gelatin, 0.5 g mL^-1^ Chromium(III) potassium sulfate; sterile filtered) for 3-minute intervals, followed by 30 seconds of gentle dipping. This process was repeated for a total of 3 cycles. Coated slides were then covered with aluminum foil to prevent debris accumulation and dried for 72 hours in a ventilated fume hood. Post-drying, slides were transferred to clean, dry slide boxes and stored at RT until use.

All tissue samples presented herein were sectioned on a cryostat (Leica). Prior to sectioning, tissue samples were transferred to a –20 °C freezer for 48 hours to ensure a homogeneous temperature. Sections were then mounted on a metal puck with excess O.C.T. and sectioned into 20-µm thick slices. Because of the relatively large volume of the injection site, tissue samples were collected at semi-regular intervals with a target spacing of 120 µm between cuts and a total of ∼45 tissue sections collected from each heart. Sections were collected on 15 pre-labeled slides, with 2 – 3 tissue sections per slide. The number of sections discarded between tissue slices were carefully recorded in spread sheets for each sample to ensure that tissue-to-tissue variability in cutting could be accounted for in final tallies (see *Microsphere and Gel-Retention Quantification*). After sectioning, any remaining tissue was discarded, and slides with heart sections were stored in slide boxes at –80 °C until mounting and imaging.

### 2.18 Injection Scoring

All injections were recorded and scored to assess the variability between cohorts of animals and subjective human interpretation of what constitutes a successful injection in comparison to immunohistochemical data. We devised a 4-part scoring system that assessed clinically relevant concerns in cardiothoracic surgical injection paradigms: (1) the observable amount of material reflux (on a scale of 0-3 points ranging from no visibly remaining material to no material reflux), (2) ease of injection (on a scale of 0-3 points ranging from no smoothness in injection to controlled smooth injection), (3) degree of bleeding post-injection (on a scale of 0-3 points ranging from excessive bleeding with risk of death to no spot bleeding), and (4) correct location of the injection site (on a binary scale of 0-1 point with the injection visually being in the incorrect or correct location). These scores were tabulated for each gel group at both time points for a total possible score of 10 points. Scores were assigned by an external observer, blind to the injection group and time point, by reviewing the videos of each injection and giving scores for each injection based on the previously agreed upon criterion.

### 2.19 Tissue Imaging

To ensure an approximately equal number of sections were imaged for microsphere and hydrogel retention, 8 slides with 2 – 3 semi-regularly sectioned tissue slices were selected from the 15 total slides collected per animal. Slides were first removed from –80 °C storage and then warmed to RT in a shadow box to avoid photobleaching of fluorescent microspheres or embedded fluorescently labeled ELP. After warming, slides were then incubated at 37 °C for 30 minutes to melt the surrounding O.C.T. and subsequently rinsed in tap water for 5 minutes to clear O.C.T. from the slide and tissue. After removal of the O.C.T., tissue sections were carefully dried of excess water on the back and sides. With careful pipetting to avoid creating bubbles, 40 µL of ProLong anti-fade media (Cell Signaling, 9071S) and a coverslip (Thermo, 3323) was then used for mounting and sealing. Slides were left to dry at room temperature in a shadow box for 72 hours, and then imaged.

Fluorescent imaging of heart sections was performed on a Leica Thunder Inverted microscope. Tissue sections from selected slides were quickly surveyed with a 2.5X air objective, and any regions containing fluorescent microspheres or labeled ELP were then imaged with a 10X air objective. Images in the present manuscript were taken as tile scans that varied in physical dimensions depending on the size of the visible cross-section of ELP/microspheres in each tissue slice, but on average had a physical height of ∼60 µm (3.80-µm step size). Tile-scans were stitched together (20% overlap) within the Leica software (LASX) and were additionally clarified using the Leica Thunder & Lightening computational clearing algorithm to remove background noise. Importantly, all images were processed with identical clearing settings. All images are presented as max projections, compiled, and exported in

ImageJ (ImageJ opensource software) [44]. For consistency of presentation, fluorescent images were placed onto a black, square background of identical size in Photoshop and presented as representative images.

### 2.20 Microsphere and Gel-Retention Quantification

Quantification was carried out in three steps: (1) image analysis in Fiji (ImageJ opensource software) [44] (2) post-processing data with an automated Python script and (3) final tabulations in Excel (Windows). First, composite images were split into individual channels and saved as separate images to isolate microspheres and fluorescently labeled ELP for quantification. Through ImageJ macro scripting, we batch converted all images to binary images and using a particle counter script in Fiji (ImageJ opensource software) [44, 45], quantified the number of microspheres or, where applicable, the cross-sectional area of ELP. Data was recorded both as a summary count (Summary microsphere count/ELP cross-section per image) and individual data sets (full list of microspheres/ELP with corresponding cross-sectional area of each particle) for each image. In some cases where microspheres had clustered together the automated particle counting would underestimate the number of microspheres in an image. To account for this, we wrote a secondary python script that took the individual image data for each image and divided the calculated area (px²) of every particle by the average cross-sectional area of a microsphere (∼ 71 px²). Based on these values, we were able to estimate the number of microspheres that were undercounted in each image from clustering and then added these values to the respective image count to obtain a more accurate estimate.

Based on the microsphere count and ELP-cross-sectional area calculated from each tissue slice, we estimated the total number of microspheres and, for HELP gels, the estimated volume of the hydrogel. To tabulate the number of microspheres within the injection site, we summed two values over the total number of images (*N*): (1) the total microsphere count from each individual image (Bi) and (2) an estimation of the microsphere count between two image slices (*B*_*int*_) that were discarded (**Eq. 2**). To estimate the number of microspheres between two sequential tissue slices, we assumed that there was a linear transition in the microsphere count between them. Stated another way, we assumed that the slices between two collected tissue slices, ‘*i*’ and ‘*i* + 1’, had values that linearly transitioned between the microsphere count on slice ‘*i*’ (*B*_*i*_) and slice ‘*i* + 1’ (*B*_*i* + 1_). Then, by knowing the number of tissue slices that were discarded between two collected sections (*n*_*i* + 1_), we could estimate the number of microspheres in each slice and sum them (*B*_*int*_). The general form of this equation has been provided (**Eq. 3**) and is what was used in the present manuscript. For the volume tabulation, we first approximated the cross-sectional area calculated in ImageJ to a circular cross-section and determined a radius for each slice (e.g. *r*_*i*_ and *r*_*i* + 1_) (**Eq. 4**). Then, by knowing the number of tissue sections (*n*_*i* = 1_) collected between each slice and the tissue thickness (*C*), we can use the general equation for the volume of a conical section to estimate volume between them and sum them over all of our collected tissues (**Eq. 5**).

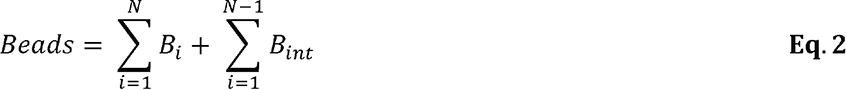

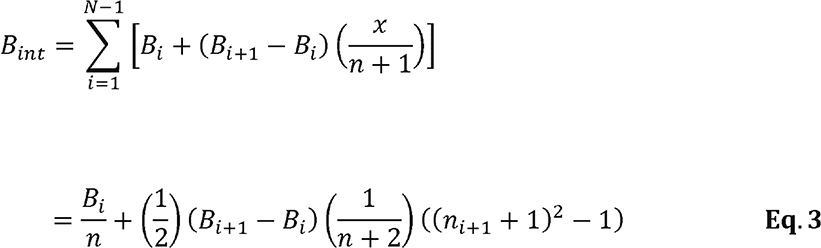

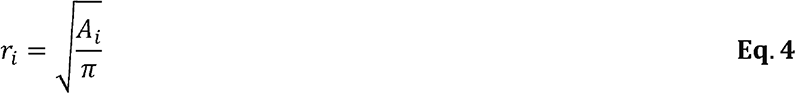

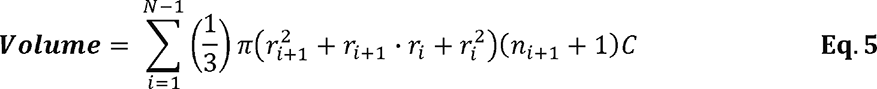

### 2.21 Hematoxylin and Eosin Staining

Heart tissue was fixed and sectioned as previously described. From each animal, a single representative slide with 2 – 3, 20-µm, thick heart sections were removed from –80 °C storage. Slides were then warmed to RT in a shadow box to avoid photobleaching of fluorescent microspheres or embedded fluorescently labeled ELP’s. After warming, slides were then incubated at 37 °C for 30 minutes to melt the surrounding O.C.T. and subsequently rinsed in tap water for 5 minutes to clear O.C.T. from the slide and tissue. After removal of the O.C.T., all slides were carried through the following general protocol. Stain with hematoxylin for 3 minutes; rinse with tap water for 1 minute; immerse in clarifying solution for 1 minute; rinse in 95% ethanol for 1 minute; stain with eosin for 30 seconds. After eosin staining, tissues were dehydrated through a step-down process. Briefly, 95% ethanol for 1 minute; two successive washes with 100% ethanol for 1 minute each; 2 successive washes with Xylene for 2 minutes each. After dehydration, slides were then dried, and covered with mounting media and a coverslip.

### 2.22 Data Analysis and Statistics

All data are presented as mean ± standard deviation (SD), with individual replicate values indicated by a black circle (•). Replicate count is either listed in the figure directly or in a corresponding methods section. Statistical significance testing for all data was performed in the GraphPad Prism 9 software package. As a variety of statistical tests were used, the specific test for each data has been listed in the corresponding figure cation. In general, we used a significance cutoff of α = 0.05 and a relevant post-hoc test that is also included with the description of each statistical test. In all figures, significance cutoffs are defined as follows: * p < 0.05; ** p < 0.01, *** p < 0.001, **** p < 0.0001, or n.s. for not significant.

## 3. Results and Discussion

### 3.1 HELP variant synthesis and mechanical characterization

Our work was motivated by the clinical need for materials with mechanical properties that both allow catheter injectability and significant retention within the contracting myocardium. Due to their low viscosity, saline or medium are commonly used as a catheter-injectable delivery systems, yet their ability to retain therapeutics within the myocardium is low [46–51]. A previously reported variant of our hyaluronan and elastin-like protein (HELP) hydrogel system was used to deliver cells and miRNA to the heart in preclinical studies [52], yet no quantitative measurements of retention were performed, inhibiting optimization of gel mechanics for clinical translation. Similarly, another requirement for clinical translation is surgical feasibility, which requires an exploration of the correlation between gel mechanics and injectability. To address these questions, we chose to synthesize a family of five HELP hydrogel variants with distinct mechanical properties for characterization of injectability.

Our HELP gel system is a dynamic covalently crosslinked (DCC) polymer network. Upon mixing, the hydrazine-functionalized elastin-like protein (ELP) component forms a DCC hydrazone linkage with either an aldehyde- or benzaldehyde-functionalized hyaluronan (also called hyaluronic acid; HA). The recombinant ELP (full amino acid sequence in **Figure S1**) was expressed in *Escherichia coli*, purified through temperature cycling, and conjugated with hydrazine moieties as previously published [40]. Two different synthetic routes for preparation of the HA component were performed to compare their effects on overall HELP gel mechanical properties and injectability. First, a commercially available, recombinant HA (M_w_ ∼ 1.5 MDa) was oxidized using sodium periodate exposure for 8, 16, or 24 hours; creating our first three variants termed HA8, HA16, and HA24, respectively. Alternatively, a commercially available, recombinant HA (M_w_ ∼ 100 kDa) was functionalized through bio-conjugation such that 12% of the HA backbone structure was modified with a pendent alkyne group (HA-Alkyne). Then, the resulting HA-Alkyne was subsequently modified with a copper-click reaction to conjugate either aldehyde groups (HA-A) or benzaldehyde groups (HA-B) (**Figure S2**).

To probe how these different modification schemes influence gel mechanics, we kept the overall gel concentration the same with a final 2% (w/v) ELP and 1% (w/v) HA, and only altered the identity of the HA component between our five variants. Using oscillatory shear rheology, we found that all five formulations showed evidence of gelation (storage modulus (G’) > loss modulus (G’’)) in under 30 seconds (onset of gelation occurred before beginning the test for all samples) (**Figure S3**). Additionally, after 30 minutes of crosslinking at room temperature (RT), all five gels had reached a stable plateau modulus indicating complete crosslinking (**Figure S3**). Following a heating step to 37 °C, which induces a secondary physical crosslinking step due to self-assembly of the elastin-like peptides [37], we tabulated the average shear modulus (**Figure 2A**) and stress-relaxation rate (**Figure 2B**) from 3 – 5 independent replicates and found that both properties can be significantly changed by altering the identity of just the HA component. For reference, we present the tabulated averages (**Figure 2C**), representative frequency sweeps (**Figure 2D**), and normalized stress-relaxation curves (**Figure 2E**).

**Figure 2.**
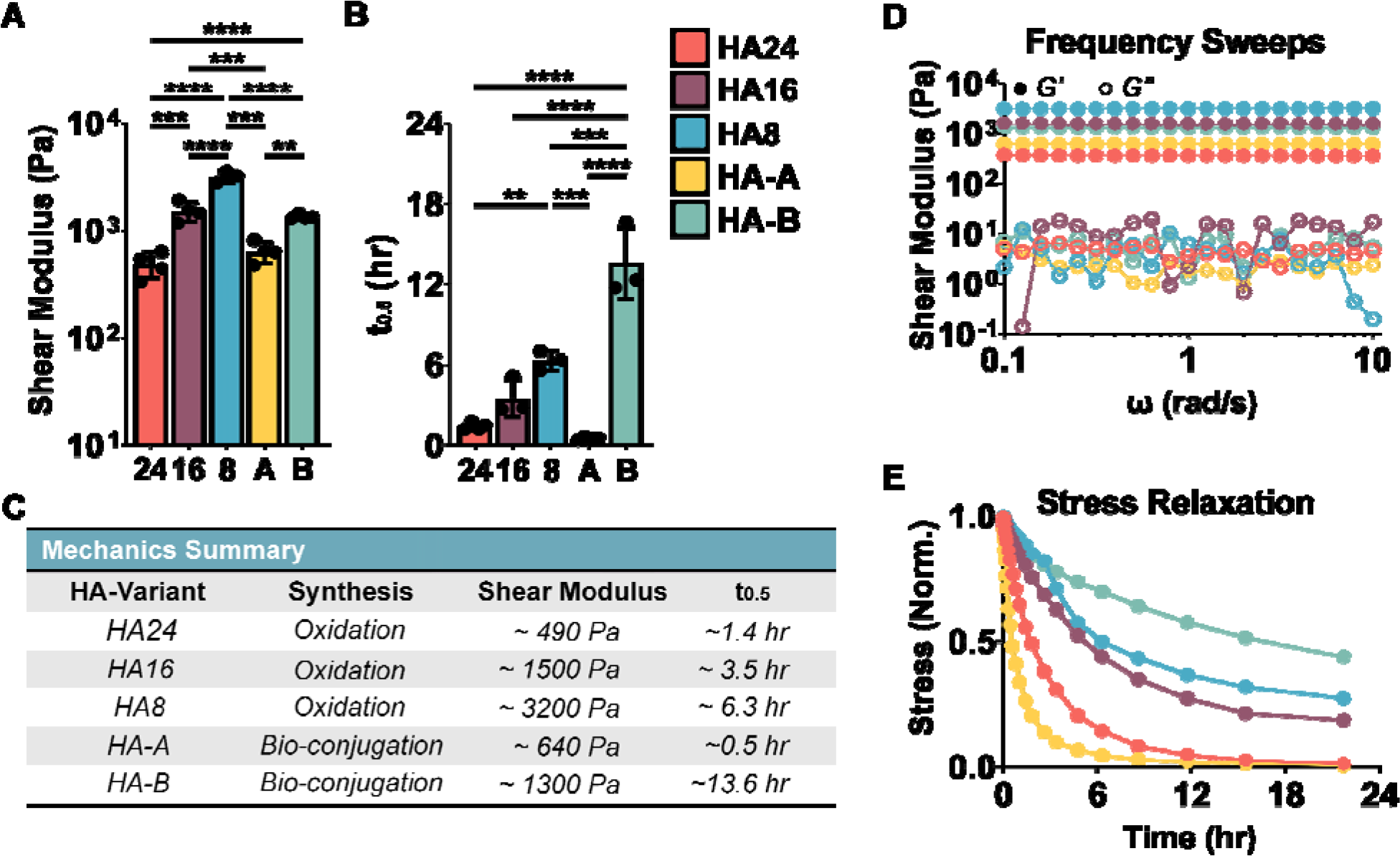
Hyaluronic acid and Elastin-Like Protein (HELP) Hydrogel Mechanics. Five HELP gels (all 2% (w/v) ELP and 1% (w/v) HA) were synthesized; three variants used oxidation for 24, 16, or 8 hrs (HA24, HA16, HA8, respectively); two variants used bio-conjugation chemistry to append aldehyde or benzaldehyde groups onto HA (HA-A, HA-B, respectively). (A) Summary shear rheology frequency sweep and (B) stress relaxation data for our five HELP gels. (C) Average values of maximum storage moduli and stress relaxation time (t_0.5_) for each gel type (n = 3 – 5) have been summarized in table. Representative (D) frequency sweep and (E) normalized stress-relaxation curves are additionally presented for reference. The HELP system forms robust gels with a shear modulus range from ∼490 Pa to ∼3200 Pa and range of stress relaxation times from ∼0.51 hr. to 13.6 hr. One-way ANOVA, α = 0.05, post-hoc Tukey test, ** p < 0.01, *** p < 0.001, **** p < 0.0001.

These data indicate that by changing the identity of the HA component we can modulate the resulting shear modulus between ∼490 Pa and ∼3200 Pa while the stress-relaxation rate, quantified as the time to dissipate half of the initial stress at a constant strain (10%), ranged between ∼0.51 hr to ∼13.6 hr. For the gels functionalized through oxidation, decreasing the oxidation time resulted in stiffer gels, *i.e. G’* for HA24 < HA16 < HA8. Similarly, decreasing the oxidation time resulted in gels with longer stress relaxation times, *i.e.* t_0.5_ for HA24 < HA16 < HA8. When comparing the two bio-conjugated formulations, HA-A *vs*. HA-B, we find that the stress-relaxation rate is dependent on the exchange kinetics of the hydrazone bond, with aldehydes having a faster exchange rate than benzaldehyde and, therefore, a faster stress-relaxation time, consistent with other reported literature [39, 53]. For context, our measured shear stiffness moduli are less than what has been reported for cardiac tissue, 5 – 50 kPa [54, 55]; however, direct comparisons are difficult due to variations in testing methodology. As another comparison point, Matrigel, a common injectable hydrogel for preclinical studies, has a measured shear stiffness modulus that is two orders of magnitude lower, ∼39 Pa (**Figure S4**). Similarly, while stress-relaxation properties have been characterized for several tissues in the body [56], sources for cardiac tissue are somewhat limited. One study using a biaxial and uniaxial stress-relaxation test on aortic valve leaflets found that relaxation with a maximum membrane tension of 60 N m^-1^ was directionally dependent, with the circumferential and radial directions relaxing ∼27.5% and ∼33.28%, respectively, in three hours [57]. When using our testing protocol to characterize Matrigel, a relatively fast stress-relaxation rate of 67 sec was observed (**Figure S4**).

### 3.2 Qualitative evaluation of in vitro injectability

Following mechanical characterization, our five HELP formulations were screened for their qualitative injectability. DCC polymer networks based on hydrazone linkages have been shown by our group and others to be shear thinning and injectable [37]. We designated three criterion that we deemed vital to facilitate repeatable injections (**Figure 3A**). First, to facilitate clinical translation, we required that (1) gels should be fully crosslinked prior to injection. While many injectable gel formulations are designed to crosslink *in situ*, this often results in a narrow time window within which the gel can be injected. If the *in situ* crosslinking is delayed, the material can flow away from the target injection site or be ejected out of the contracting myocardium. In contrast, if the surgeon injects too late or too slowly, then premature crosslinking can result in a clogged needle and failed delivery. Designing a gel to be crosslinked prior to injection overcomes these limitations. Secondly, we required that (2) gels had to be able to be easily injected through a 30-G needle with hand force by an experienced animal surgeon. Lastly, to reduce the possibility of trauma to the surrounding tissue that may cause excessive bleeding, we required that (3) gels had to have a smooth injection with no evidence of a ‘burst’ injection. To mimic the protocol for preclinical injection studies, 50 µL of HELP gel at a final concentration of 2% (w/v) ELP and 1% (w/v) HA was made for each HA-variant and injections were repeated at least three times to test for consistency. Full videos of a representative injection for each variant can be found in **Figure S4**. Individual stills have been provided in **Figure 3B**.

**Figure 3.**
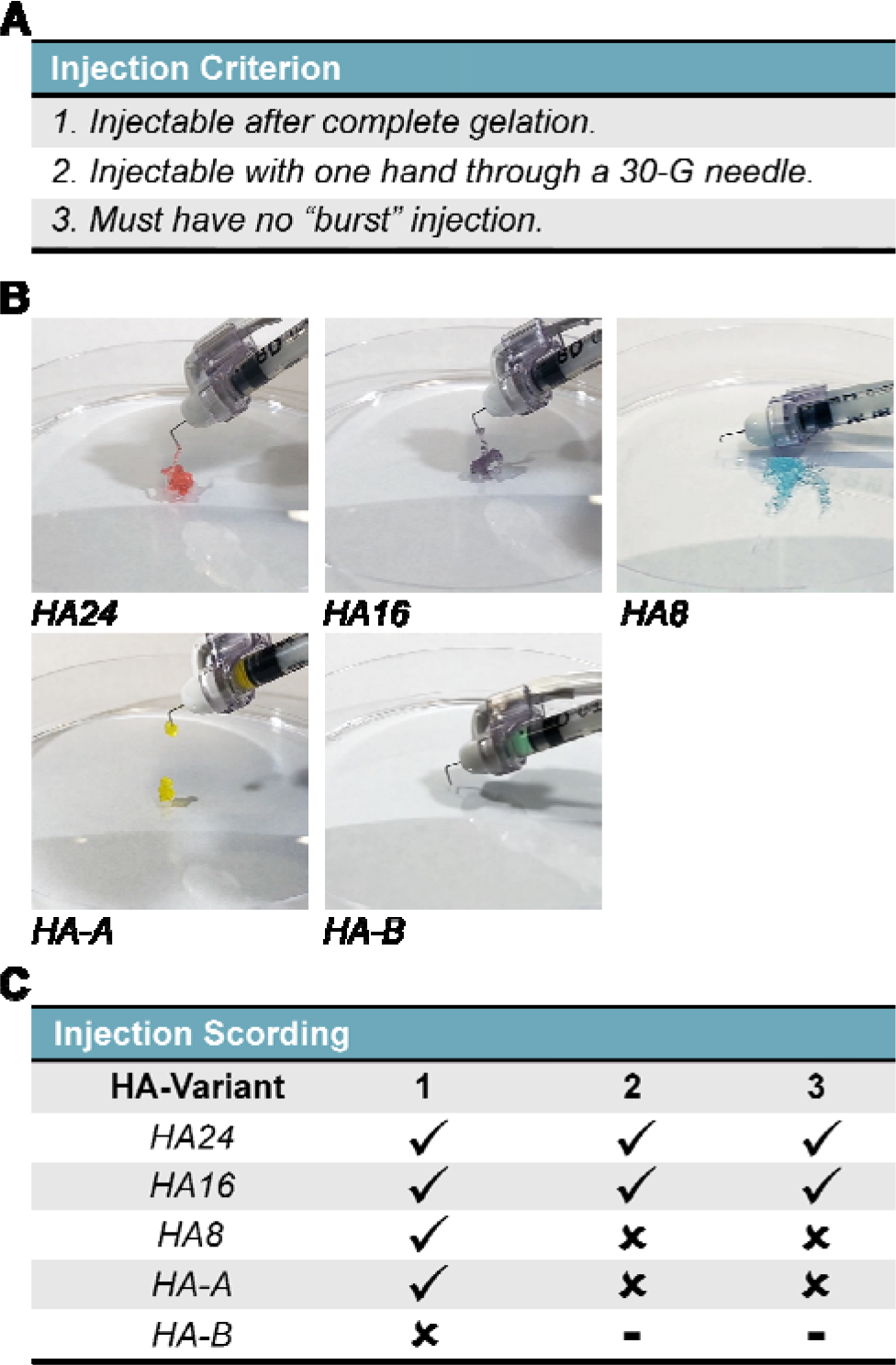
Hydrogel Injectability Screening and Qualitative Assessment. The qualitative injectability of HELP gels was tested by injecting each formulation through a hooked, 30-G insulin needle. (A) Injectability criteria included: (1) ability to be injectable after complete gelation (30 min), (2) injectable with one hand, and (3) no evidence of a sudden burst injection. (B) Qualitative stills taken from recordings of each injection. Note: brightness has been artificially increased to increase ease of viewing. Additionally, each gel has been dyed with food coloring to match their prescribed color scheme as presented in this manuscript. (C) Table summarizing the qualitative assessment, where a check mark (?) and a cross (x) signify passing or failing the test, respectively. HA-B was not injectable at all, and so criterions (2) and (3) could not be evaluated and have been designated with a ( - ).

The results of our assessment are summarized in **Figure 3C**, where a check mark (?) and a cross (x) signify passing or failing the test, respectively. We found that only two of our five HA-variants, HA24 and HA16, created HELP gels that met all three criteria. The HA-8 and HA-A variants could be injected post-crosslinking, but required the surgeon to use both hands to apply an adequate amount of force. In our experience, stabilization of the injecting hand is required to limit tissue damage and ensure a consistent injection force, and so gels that require the use of both hands pose increased risk to the animal and a higher degree of variability. We additionally found that both HA8 and HA-A had evidence of a ‘burst’ injection where all (HA8) or a large fraction (HA-A) of the material shot out of the needle during injection. We postulate that a “burst” injection could cause the breakage of local vasculature and potential damage to the surrounding tissue, which could adversely impact gel and payload retention. The HA modified with a benzaldehyde group, HA-B, could not be injected despite the operator using both hands.

Based on the encouraging results of our injectability study with a 30-G needle (0.8 cm long, 0.159 mm inner diameter), we wanted to further verify the potential for translation using clinical delivery systems. As a proof-of-concept demonstration, we injected 700 µL of HA16 (dyed green for ease of visibility) through a 150-cm medical catheter (Codman, 606-151MX, 0.75 mm inner diameter tube, 0.65 mm inner diameter end nozzle) (**Figure S6**). While a greater amount of hand force was needed to move the material through the length of the catheter compared to the shorter, 30-G needles, gel was successfully and smoothly ejected with a steady flow from the tip of the catheter (*i.e.* no “burst” injection), suggesting to us that catheter delivery of HELP gels is possible and could potentially be implanted into larger mammals via intravenous delivery. This is a particularly important step to validate, as catheter injectability post-crosslinking is vital to side-stepping the need for thoracic-level surgeries for implantation.

### 3.3 Quantitative assessment of gel failure stress and yield stress

Following *our in vitro* injectability study, we sought to characterize which mechanical properties of our five hydrogel systems best served as a ‘predictor’ for injectability. Based on our initial mechanical characterization, there did not appear to be a direct relationship between either the initial matrix stiffness nor the stress-relaxation time (as indicated by t_0.5_) (**Figure 1C**) and ease of injection. HA-A, for example, was both ‘softer’ (*i.e.* more compliant) and had a shorter stress-relaxation time than HA16, but was markedly harder to inject through a 30-G needle. Given that the injection occurs post complete cross-linking, and on a time scale that is much faster than the stress-relaxation time of our hydrogels, we hypothesized that the injection may be plastically deforming the hydrogel through a mix of bond and chain breaking, as has been suggested for other dynamic hydrogels [58–60].

To test this hypothesis, we ran a two-part recovery experiment. First, we injected 50 µL of our two injectable gels, HA24 and HA16, directly onto the rheometer post-crosslinking through a 30-G needle and monitored changes in mechanical properties over the course of 24 hours at a low strain (1%) to test if the network would self-heal and recover their initial mechanical properties. Then, we ran a high strain (10%) post-recovery frequency sweep, identical to those in **Figure 2**, to see if the mechanics at high strain had been recovered. Our measurements show that the gels rapidly self-heal back into the gel-phase after extrusion, with *G’ > G”* before the sample measurement could begin (**Figure S7A, B**). Over the next hour, there is a slight, gradual increase in the shear moduli for both HA16 and HA24, suggesting further self-healing, perhaps through the reformation of dynamic hydrazone bonds. Both gels reach an approximate plateau shear modulus of ∼300 Pa at 24 hours post-injection. Notably, this plateau shear modulus is lower than the pre-injection plateau modulus; ∼450 Pa and ∼730 Pa for HA24 and HA16, respectively (**Figure S3**), suggesting that full network recovery is not achieved. Consistent with these data, our post-recovery high strain test (10%) shows that both HA16 and HA24 cannot sustain this high level of deformation and enter the sol phase (*G” > G’,* **Figure S8A, B**, respectively). These data suggest that the network has undergone some amount of plastic deformation relative to the initial gel network structure, which can sustain these same high strains (**Figure 2D**).

We hypothesize that upon flow, these gels fracture into discrete gel “blobs” that then flow as discrete microgels with “plug-flow” fluid dynamics [61]. Other dynamic hydrogels, including those formed through peptide and protein self-assembly mechanisms appear to have similar flow mechanisms [58–60]. We hypothesize that once the gel is ejected, the system of discrete microgels rapidly recovers a macroscopic, bulk gel phase due to dense microgel packing (sometimes termed a “granular hydrogel” [62, 63]). Over time, dynamic covalent hydrazone bonds can reform between adjacent microgels to further strengthen the recovered hydrogel.

To probe this theory further, we ran a series of successive creep tests at progressively higher stress (*σ*) states and measured the responsive strain (γ) of our hydrogels (**Figure S9**).

Importantly, after each creep test we included a relaxation period ( for 1,000 sec) to allow for recovery of elastic deformation. From these data, we calculated a strain rate (*dγ*/*dt* = *γ̇* ) for each stress state, and by applying Newton’s law of viscosity (**Eq. 1**), we can report an apparent viscosity (*η*) for each stress state. Given that the apparent viscosity is linear within the elastic regime (*η* = *σ/γ̇*), we estimated the apparent yield stress (σ_*y*_) and fracture stress (σ_*F*_) by monitoring for deviation from linearity at each applied stress. In the present work, we define σ_*y*_ as the stress-step that induces a permanent deformation (% γ » after the relaxation period) and σ_*F*_ as the stress-step that induces failure of the gel (η → 0 ).

Representative plots of apparent viscosity measurements for our successive creep tests have been reproduced in **Figure 4A**. For all five HELP gels, the apparent viscosity calculated across successive stress steps was relatively constant (**Figure 4A**), confirming that our testing parameters were appropriate for this set of materials. For HELP gels produced through oxidation chemistry (HA24, HA16, HA8), we did not observe yielding behavior across this range of applied stresses, while HELP gels produced through conjugation reactions (HA-A and HA-B) had clear evidence of yielding (**Figure S9**). Multiple, independent replicates (n = 4 – 5) of each HELP gel variant were subjected to the successive creep test to characterize the average fractures stress (**Figure 4B**), which was tabulated for all gels (**Figure 4C**). The observed fracture stress ranged from approximately ∼170 Pa to ∼12,000 Pa, following HA24 < HA16 < HA8 < HA-A < HA-B. These data parallel our qualitative *in vitro* screening of injectability that had an identical ordering of relative difficulty of injection (**Figure 3**). Taken together with the rheological data from post-injection gels (**Figure S8**), these results suggest that the injectability of our HELP gels is made possible through plastic deformation of the gel network and further supports the use of *σ*_*F*_ as an indicator of relative injectability for this system.

**Figure 4.**
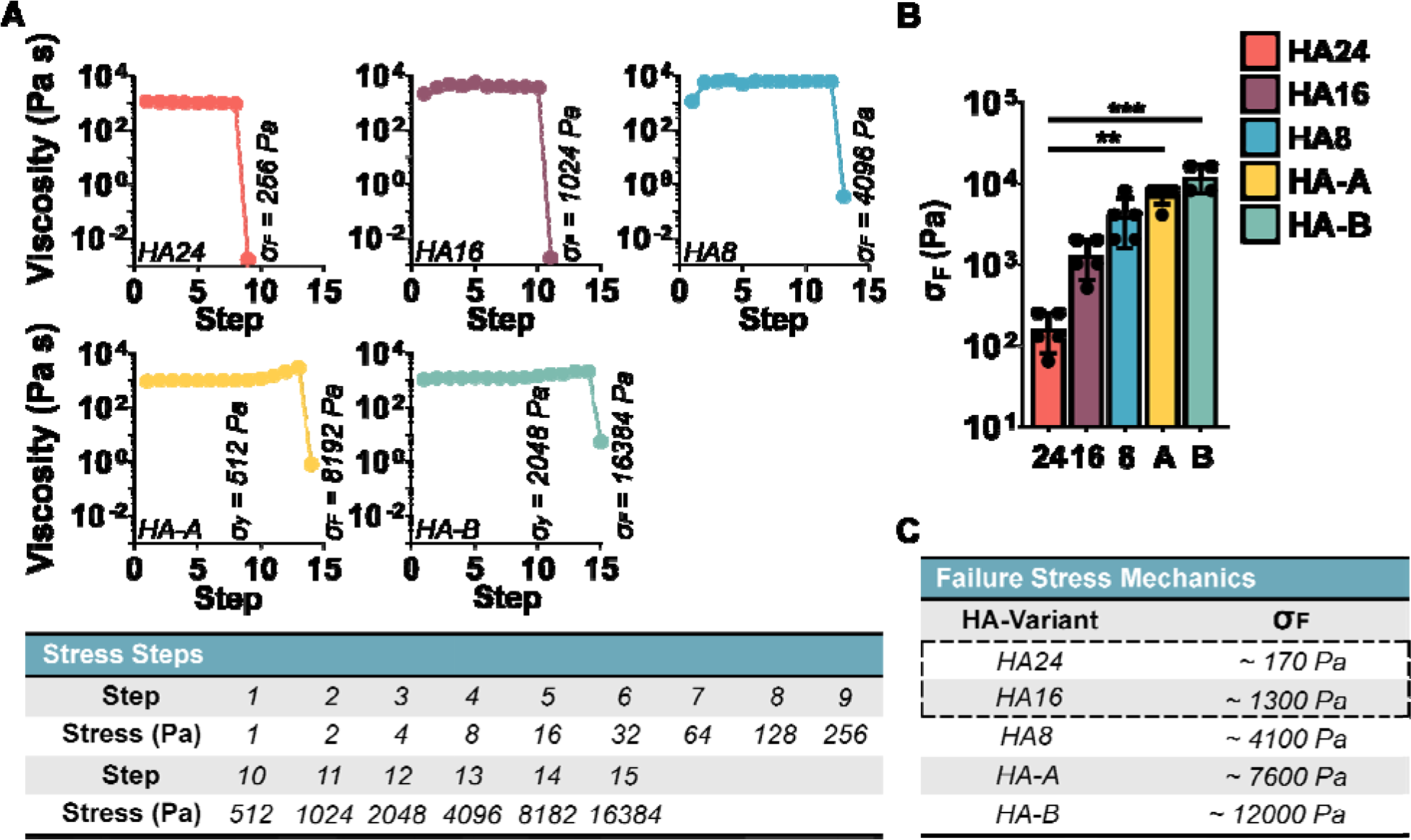
Hydrogel Fracture Stress. To quantify fracture stress ( ) we measured the gel apparent viscosity ( ) as a function of stress, with increasing stress intervals. The gel fracture stress ( ) was defined as the stress step that led to an abrupt decrease in viscosity. (A) Representative measurements of apparent viscosity at increasing stress step intervals. denotes failure stress;denotes yield stress. (B) Average fracture stress for each HELP gel formulation (n = 4 – 5). (C) Tabulated fracture stress values demonstrate that the two injectable formulations identified in Figure 2 (HA24 and HA16, designated with a dotted outline) have the lowest fracture stress. Kruskal-Wallis test α = 0.05, with post-hoc Dunn’s test, ** p < 0.01, *** p < 0.001.

It is interesting to consider the broad range of failure stresses displayed by our materials despite having identical compositions. One likely source of this variation in failure stress is from the differing molecular weights within our samples. The oxidation of HA is known to significantly reduce the molecular weight, which can in turn influence the injectability [60]. Using gel permeation chromatography (GPC) we characterized the molecular weight of our experimental

HA groups as well as an array of commercially available, unmodified HA samples (M_W_ range of 20 kDa to 1.5 MDa) as controls (**Figure S9A - C**). Consistent with the work of others, we found that the molecular weight of our oxidized variants significantly decreased with oxidation time, with the initial M_W_ of 1.5 MDa dropping to apparent M_w_ of 177 kDa, 80.1 kDa, and 56.8 kDa for HA8, HA16, and HA24, respectively.

In contrast, the HA variants synthesized by conjugation chemistry had negligible changes in observed molecular weight: the initial M_W_ of 100 kDa resulted in final apparent M_W_ of 221 kDa and 141 kDa for HA-A and HA-B, respectively, as observed by GPC. These observed molecular weights are somewhat larger than the theoretically predicted M_W_ for a 12% HA modification reaction, which are 115 kDa and 105 kDa for HA-A and HA-B, respectively. This discrepancy is likely due to the bulky and hydrophilic side-chains causing the HA polymer chain to expand, which was confirmed by dynamic light scattering wherein we measured the hydrodynamic radius (R_h_). The measured R_h_ of all HA molecules used in this work, including unmodified controls of varying M_W_; the oxidized HA8, HA16, and HA24; and the conjugated HA-A and HA-B molecules along with the intermediate HA-alkyne are all shown in supplemental data (**Figure S10D - F**). Consistent with our GPC observations, unmodified, 100 kDa HA and 12% modified HA-Alkyne had a measured R_h_ of 10.56 nm and 10.95 nm, respectively, while HA-A and HA-B had R_h_ of 14.48 nm and 11.51 nm. These data suggest that the addition of -Ald and -Bza side-groups led to an increase in R_h_ which led to an over-prediction of the observed M_w_ by GPC.

Taken together, our data suggest that gel fracture stress plays a significant role in influencing the injectability of the gel. Other groups, particularly in the field of 3D bioprinting, have hypothesized that the extrusion of hydrogels with DCC hydrazone linkages may have a component of brittle fracture that enables them to be extruded [60]. Our data are consistent with this idea, with there being little to no yielding behavior prior to the abrupt failure of the material ( 0). Our data further demonstrate that both molecular weight and hydrazone bond kinetics can influence σ_*F*_. For our oxidized gels (HA8, HA16, and HA24), we observe that gels with lower molecular weight, and hence fewer chain entanglements, display lower fracture stress and easier injectability (**Figures S9, 3, 4**). For our bio-conjugated gels (HA-A and HA-B), the formulation with faster hydrazone bond kinetics, and hence faster stress relaxation rate, displays lower fracture stress and easier injectability (**Figures 2 - 4**). Lastly, we can conclude that HELP gels that have _\:’_ values < ∼1300 Pa (HA16 and HA24) are injectable post-crosslinking (**Figures 3 - 4**).

These general trends will be useful for the design of future injectable gels. These data also suggest that more thorough analysis across a range of bond types, average chain length, and variation in chain length (poly-dispersity index; PDI) may assist in uncovering the interacting influences between these variables and, ultimately, the resulting failure stress ( _\:’_). Lastly, for these specific studies, we determined that HA16 and HA24 would be the most viable candidates for clinical translation, and hence these materials were further evaluated within an *in vivo* retention model.

### 3.4 In vivo preclinical retention study

To test the *in vivo* efficacy of our injectable HELP variants (HA24 and HA16), we quantified the relative retention of 10-µm fluorescent microspheres injected directly into healthy, beating rat myocardium both 1 day and 7 days post-injection. The choice of using a healthy, rather than a diseased or injured, murine model was made (1) to increase reproducibility (since common injury models like complete ligation of the left anterior descending coronary artery typically results in a range of infarct lesion sizes [64]), and (2) because retention of cargo is significantly more challenging in beating tissue compared to non-beating tissue [28, 29]. Matrigel was selected as a commercially available comparison material because it is frequently used as a cardiac delivery vehicle in preclinical models [32,65,66].

For cargo quantification, all gel samples (HA16, HA24, and Matrigel) were loaded with the same concentration of fluorescent beads, 1.25 mg mL^-1^, to allow cross-comparisons.

Additionally, for HELP gels (HA16 and HA24), 20% (v/v) of the ELP peptides were labeled with a fluorescent cyanine-5 group to allow for direct imaging of the gel, as described in the methods. These concentrations and degree of labeling were determined through *ex vivo* screening to identify values that would enable automated image quantification (**Figure S11**). For our animal groups, 10-week-old Sprague Dawley rats were randomly divided into three, sex-matched tests groups: 1) HA16, 2) HA24, and 3) Matrigel, each with two cohorts for eventual end-point evaluation on either 1 day (Cohort 1) or 7 days (Cohort 2) post-injection (**Table 1**). A summary of mechanical properties (shear modulus and stress-relaxation) for our three test groups have been reproduced for reference in **Figure S12**.

**Table 1.** In vivo Test Groups. Following our injection study, both the HA16 and HA24 were selected for in vivo testing along with Matrigel as a control. For testing, 38, 10-week-old, Sprague Dawley rats were divided into sex-matched cohorts: 1 and 2, which were allocated to endpoint analyses at day 1 and day 7 post-injection, respectively.

| Test Groups |  |  |  |  |
| --- | --- | --- | --- | --- |
| Hydrogel | Cohort 1 |  | Cohort 2 |  |
|  | n | Endpoint | n | Endpoint |
| HA16 | 4 ♀ 3 ♂ | 1 Day | 3 ♀ 3 ♂ | 7 Day |
| HA24 | 3 ♀ 4 ♂ | 1 Day | 3 ♀ 3 ♂ | 7 Day |
| Matrigel | 3 ♀ 3 ♂ | 1 Day | 3 ♀ 3 ♂ | 7 Day |

L of hydrogel with 1.25 mg μmL^-1^ microspheres were injected by hand into the wall of the beating myocardium through a 30-G, manually hooked, needle. Representative surgical photos have been included in **Figure S13**. As a first point of measurement, we devised a four-part scoring system to rate the quality of our injections. The four criteria were: 1) degree of material ‘reflux’ out of the heart at the time of injection, 2) perceived ease of hand-injection into the heart by the surgeon, 3) degree of bleeding immediately following injection, and 4) accuracy of injection location within the wall of the left ventricle as determined by relative position to the left descending coronary artery. Scores were assigned based on recordings of injections by an operator blind to the hydrogel identity. Across all three hydrogels, we did not observe a significant difference in the quality of injection for either cohort 1 or 2 (**Figure S14**). Interestingly, we observed similar degrees of bleeding post-injection across all three systems, suggesting that the observable trauma to local tissue may be invariant to the identity of the gel. We hypothesize that variability in bleeding is likely more closely tied to inherent variability in the local vasculature of the animals themselves. This further emphasizes the need for gels which can hold cargo at the injection site, as it has been proposed that one of the primary modes of cargo loss may come from rupturing the local heart vasculature and which results in injectate being evacuated from the injection site out of ruptured vessels [32,67,68].

Post euthanasia, hearts were excised, sectioned, and prepared according to **Figure S15**. The microsphere count (HA16, HA24, and Matrigel) and gel volume (HA16 and HA24) were determined from quantification of confocal microscopy images using particle counters in Fiji [44, 45] (**Figure S16**). To improve the accuracy of our automated particle counts, we developed a Python script (**Figure S17**) that analyzed the readout from each image and determined the number of particles that had been excluded from counting due to microsphere clustering, as described in the methods section. Following image quantification, the approximate bead count and gel volume was determined using **Eq. 2 – 5**, as described in our methods. Representative fluorescent images of tissue sections from all three gels have been provided (**Figure 5A - F**).

**Figure 5.**
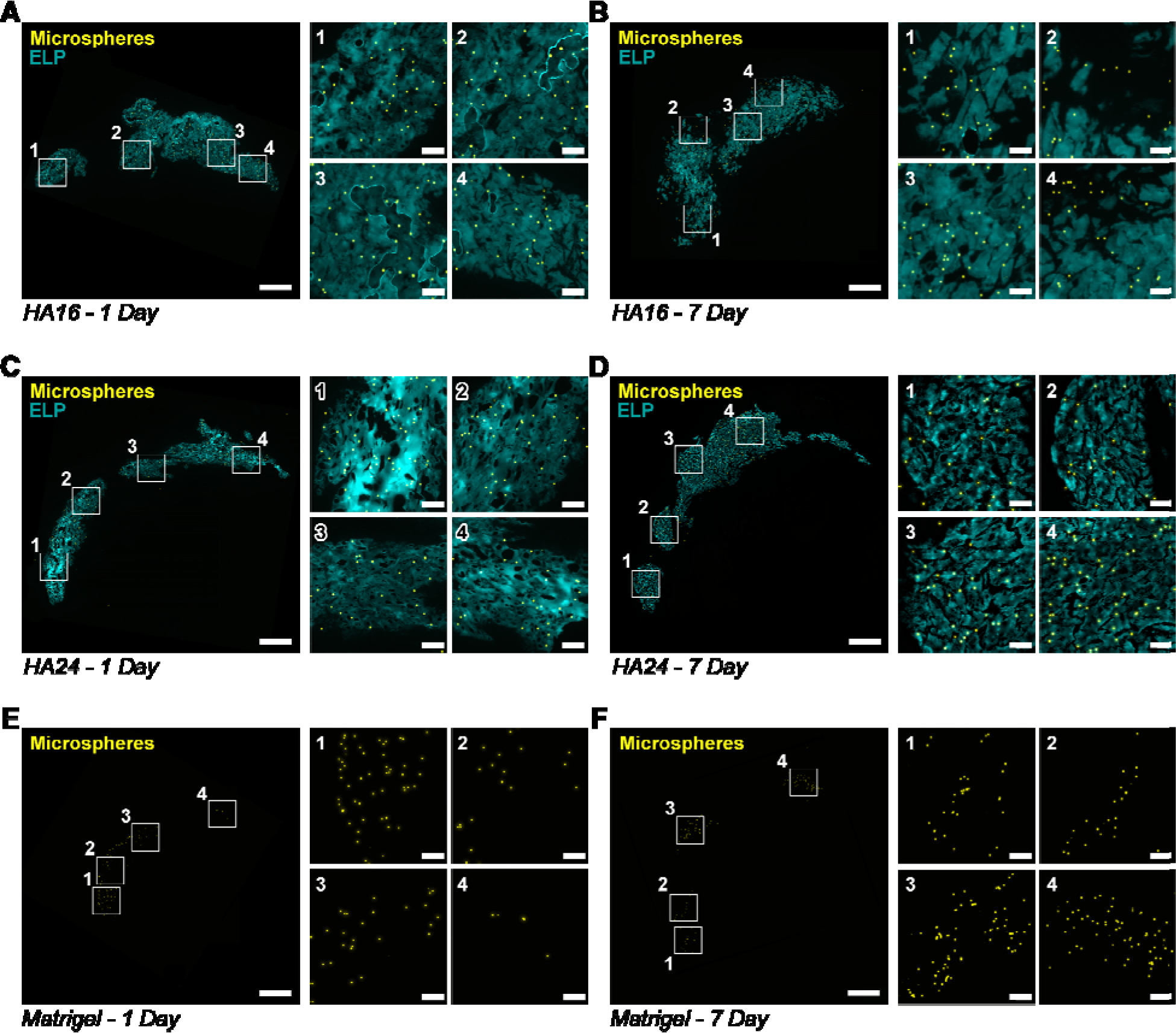
Gel-mediated cargo retention in myocardium. Representative fluorescence images (elastin-like protein (ELP), cyan; microspheres, yellow), of tissue sections collected from hearts at 1 day and 7 days post-injection with HA16 (A, B), HA24 (C, D) and Matrigel (E, F). For ease of viewing, a single low-magnification and four high-magnification (indicated by a white outline and corresponding number) fields of view are shown. Scale bar on low and high magnification images are 1-mm and 100-µm, respectively.

Qualitatively, we observed that HA16 and HA24 had a more substantial degree of spreading within the myocardium than Matrigel at both 1-day and 7-days post-injection. This was evidenced by (1) the relatively higher density of beads in Matrigel as compared to HA16 and HA24 as well as (2) the relative distance between the microspheres at the far ends of the injection region. Additionally, we observed that HA16 generally formed a more ‘bolus’ style injection, distending the tissue into a more rounded injection site than HA24, which generally spread more uniformly along the length of the left-ventricle. We note that when subdividing the animal cohorts by sex, we observed no differences in cargo retention (**Figure S18**).

Based on our quantification, HA16 significantly (* p < 0.05) improved the retention of fluorescent microspheres both 1- and 7-days post injection compared to Matrigel (**Figure 7A**). Based on our microparticle concentration, our intended cargo dosage is approximately ∼70,110 microspheres per injection (**Figure S11B**). One-day post-injection, Matrigel retained approximately 16.9% ± 12.6% of the delivered microspheres compared to 53.6% ± 24.6% and 43.1% ± 30.3% retention for HA16 and HA24, respectively (**Figure 6B, C**). Interestingly, though non-intuitively, the target bead count increased at our 7-day time point for all three hydrogels; however, these differences were not found to be statistically significant (two-way ANOVA, post-hoc Tukey test) and are likely due to experimental error.

**Figure 6.**
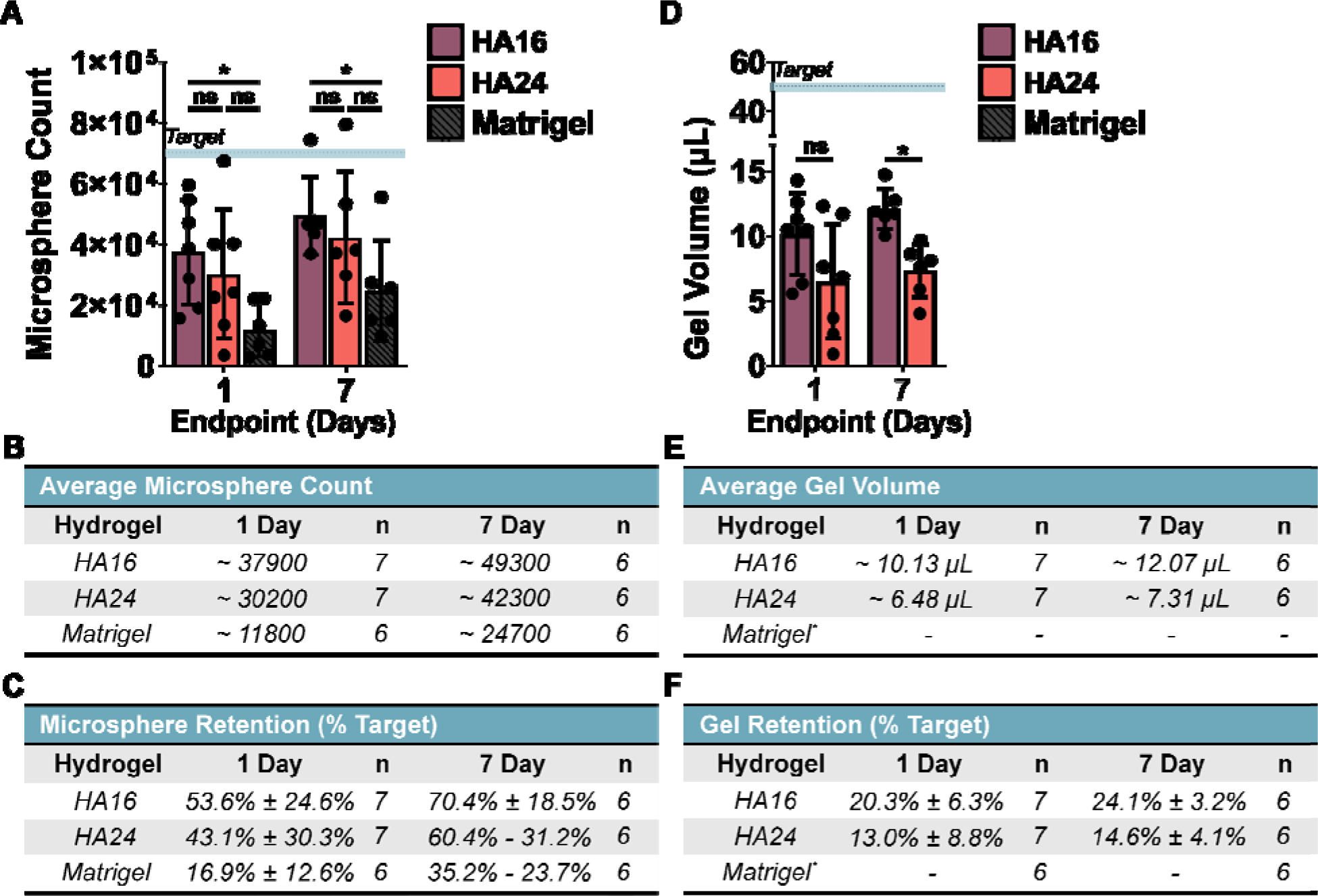
Quantified microsphere and hydrogel retention. (A) HA16 significantly increased cargo retention over Matrigel at both time points. Summary of the approximate bead count and percent retention have been provided for reference in (B) and (C), respectively. (D) HA16 had a significantly higher volume of gel retained at day 7 compared to HA24. Summary of approximate geal volume and percent retention have been provided in (E) and (F), respectively. *Note that Matrigel volume was not measured. The target microsphere count and volume are denoted by a horizontal, dotted, line corresponding to ∼70,110 microspheres and 50 µL of hydrogel. Statistical test: two-way ANOVA = 0.05, post-hoc Tukey test, * p < 0.05.

**Figure 7.**
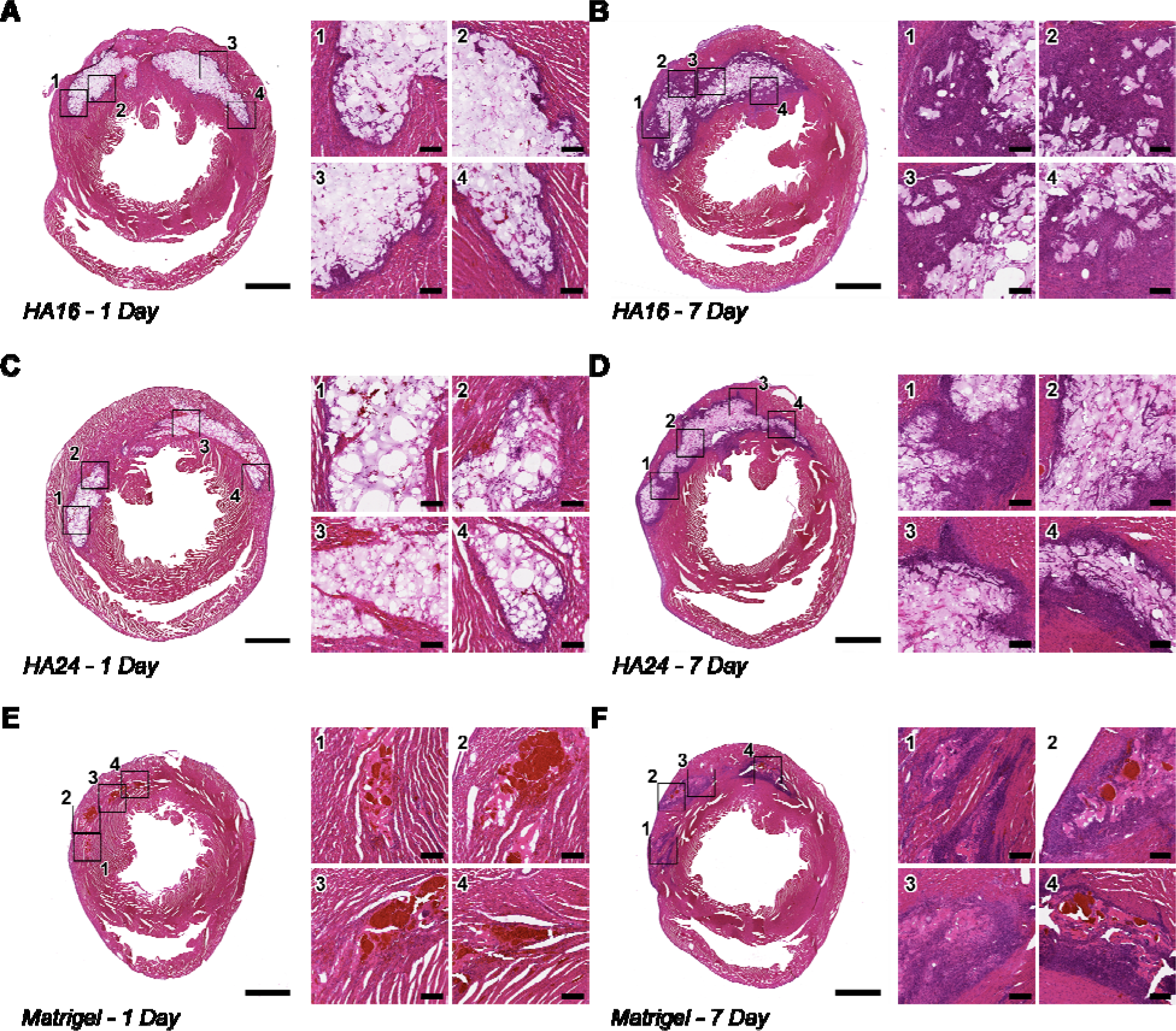
Hematoxylin and Eosin Staining of Representative Tissue Sections. Representative Hematoxylin & Eosin (H&E) stains of tissue sections from hearts 1-day (left) and 7 days (right) after injection with HA16 (A, B), HA24 (C, D) and Matrigel (E, F). For ease of viewing, a single low-magnification and four high-magnification (indicated by a black outline and corresponding numbers) have been generated. Of note, in all cases, there is an increase in cell-infiltration between Day 1 (A, C, E) and Day 7 (B, D, F) post-injection, indicated by the increase of cell nuclei (purple) in and around the implanted hydrogels. In addition, while there does not appear to be a qualitative difference in gel area between HA16 and HA24 both appear to have a greater gel cross-section than Matrigel. Note: scale bar on low magnification images represents 2-mm, scale bar on high-magnification images represents 200-µm.

The fluorescent labeling of HA16 and HA24 allowed direct quantification of the gel volume at each time point for comparison to the intended (target) gel dosage of 50 µL. Gel volume retention was found to parallel the trend in microsphere retention, with HA16 having significantly greater gel volume retention than HA24 at 7-days post-injection (* p < 0.05, **Figure 7D, E**). Similar to our microsphere counts, we saw that there was an increase between 1-day and 7-days post-injection within both groups; however, the differences between our time points were not statistically significant (two-way ANOVA, post-hoc Tukey test). Note that because Matrigel was not fluorescently labeled, an estimation of gel volume could not be made for this material. It is additionally interesting to note that the percent of gel retention is lower than the percent microsphere retention. This is likely due to underestimation of our gel-volume, stemming from our simple quantification strategy. Briefly, we ascribe a best-fit circle to describe the HELP gel cross-sectional area within each tissue slice, and then we extrapolate the volume as a conical cross-section between tissue slices. While computationally straightforward, this strategy may underestimate the total gel volume by not accounting for the complex geometry of the injection site.

As a final qualitative characterization, we stained representative sections with Hematoxylin and Eosin (H&E) **(Figure 7**). For all three hydrogels, we observed a clear increase in cell infiltration between 1- and 7-days post-injection, indicated by the increased number of nuclei (purple) within the bulk of the hydrogel. This may be evidence of an initial inflammatory response to assist with biodegradation of the injected hydrogels. We did not observe evidence of any uniform, fibrotic region surrounding the injected gels. These data encourage future detailed studies of gel biodegradation and the local immune response over time.

## 4. Conclusion

The hyaluronan and elastin-like protein (HELP) gel system presented here has several benefits compared to other currently available, injectable hydrogel systems. These include: (1) independent tuning of biochemical and biomechanical cues [38], (2) secondary crosslinking mechanisms induced at physiological temperatures [37], (3) high degree of cell-compatibility [37–39], and (4) cell membrane protection during injection [37]. Additionally, hyaluronan (HA) has been shown to promote angiogenesis, suppress fibrous tissue formation, and natively bind water to the heart muscle, which improves mechanical and electrophysiological properties of the heart muscle [69–71].

In the present manuscript, we screened a series of formulations of our two-component HELP hydrogel system for *in vitro* injectability and *in vivo* retention of microspheres following injection into the rat myocardium. Specifically, we modulated hydrogel mechanics by altering the HA component to modify molecular weight and the identity of functional side-groups to influence the kinetics of dynamic crosslink exchange. Through a series of test-injections, we found that two formulations (HA16 and HA24) could be injected easily by hand through a 30-G needle, whereas the other systems either required excessive force resulting in a ‘burst’ injection (HA8, HA-A) or could not be injected at all (HA-B). Thus, both lower polymer molecular weight and faster kinetics of dynamic crosslink exchange were found to improve gel injectability. We additionally showed that HA16 could be injected through a 150-cm catheter suggesting feasible clinical translation. In further probing the mechanics of injection, we found that injectability of our hydrogels correlated with failure stress ( _\:’_) as determined by shear rheology, suggesting that this is a useful property to characterize for future biomaterials to assist in predicting injectability of dynamic covalently crosslinked systems (DCC). Finally, we conducted an *in vivo* retention study where we looked at the retention of 10-µm microspheres when delivered with (1) HA16, (2) HA24, or (3) commercial Matrigel directly into the healthy, contracting rat myocardium. HA16 significantly improved cargo retention over Matrigel at both time points within this preclinical model.

Looking ahead at translation of these gels to a preclinical injury model, some practical considerations may present limitations. First, the rate of gelation is rapid (< 30 s), which makes homogenous mixing a challenge, especially for larger gel volumes, which would be required for any clinical-scale trials. Potential strategies to overcome this limitation include use of static mixers to permit rapid, homogenous mixing. Another point of consideration is that when working with DCC networks, the stress-relaxation effect can be used to modulate cellular response, but it could also significantly influence the stability of the gels *in vivo*. On one hand, stress-relaxation has been shown to significantly influence cell-behavior and is a valuable control point when trying to modulate cell behavior for a given application [56]. However, from the standpoint of retention post-injection, DCC polymer networks, as with all dynamic networks, are susceptible to erosion and potentially more rapid biodegradation. Thus, future translation of DCC-based networks will need to carefully evaluate the *in vivo* erosion and biodegradation rate for optimization with the intended timeline of the therapy.

In summary, the HELP gel system was found to be easily hand-injectable through both syringe needles and catheters and to significantly improve the acute retention of cargo in the contractile myocardium. These data strongly support the future investigation of these materials for delivery of therapeutic agents to challenging environments such as mechanically active tissues.

## Acknowledgements

We acknowledge N. Tran (Stanford; CA, USA) for assistance in performing the preclinical model. We thank T. Yamaoka, Y. Kanbe, and T. Takizawa (National Cerebral and Cardiovascular Center; Osaka, Japan) for preclinical model training. We acknowledge B. LeSavage (Stanford; CA, USA) for helpful conversations about statistical analyses and data presentation. We thank P. Chu and D. Wu (Stanford Human and Animal Histology Services, Stanford; CA, USA) for help with preparation of histological specimens. This work was supported by the National Institutes of Health (NIH) Training Grant in Biotechnology (T32-GM008412) to R.A.S. and R01 EB027666, R21 HL138042, and R01 HL151997 to S.C.H.

## 5. CRediT Author Statement

**Riley A. Suhar:** Conceptualization, Investigation, formal analysis, resources, writing – original draft, data curation, visualization, project administration; **Vanessa M. Doulames:** Investigation, formal analysis, resources, writing – review & editing; **Yueming Liu:** Investigation, data curation; **Meghan E. Hefferon**: Investigation, resources; **Oscar Figueroa III**: Software, validation; **Hana Buabbas:** Resources; **Sarah C. Heilshorn:** Conceptualization, writing – review & editing, supervision, funding acquisition.

## 6. Data Availability

The raw data required to reproduce these findings will be made available upon request.

## Notes

### Competing Interest Statement

The authors have declared no competing interest.

