## Supplemental Information for "Hyaluronan and Elastin-Like Protein (HELP) Gels Significantly Improve Cargo Retention in the Myocardium"

| ^a^ | Department of Materials Science and Engineering, Stanford University, Stanford, California 94305, USA |
| --- | --- |
| ^b^ | Department of Neurosurgery, Stanford University School of Medicine, Stanford, California 94305, United States |
| ^c^ | Unaffiliated |
| ^d^ | Department of Biology, Stanford University, Stanford, California, 94305, United States |
| ^*^ | Corresponding author. (S.C.H.) Address: 476 Lomita Mall, McCullough Room 246, Stanford University.   Stanford, CA, 94305-4045, USA Fax Number: 650-723-3044 |

**Summary:**

Figure S1: Elastin-like protein amino acid sequence information

Figure S2: summary of conjugation and oxidation reactions for hyaluronan

Figure S3: Representative time sweeps

Figure S4. Summary of animal mechanics

Figure S5: Representative injection videos

Figure S6: Catheter injection video of HA16

Figure S7: Recovery experiment for HA16 and HA24

Figure S8: Post-recovery frequency sweep

Figure S9: Representative stress and strain curves

Figure S10: Estimating molecular weight of modified hyaluronic acid

Figure S11: Ex-Vivo Hydrogel Fluorescence and Microsphere concentration determination

Figure S12: Summary of Animal Mechanics

Figure S13: Representative Injection Images

Figure S14: Injection Scoring

Figure S15: Tissue Sectioning Schematic

Figure S16: Example Images of Microsphere and Hydrogel Retention Quantification

Figure S17: Python Script for Post-Processing Microsphere Count

Figure S18: Sex-Based Differences in Microsphere Retention

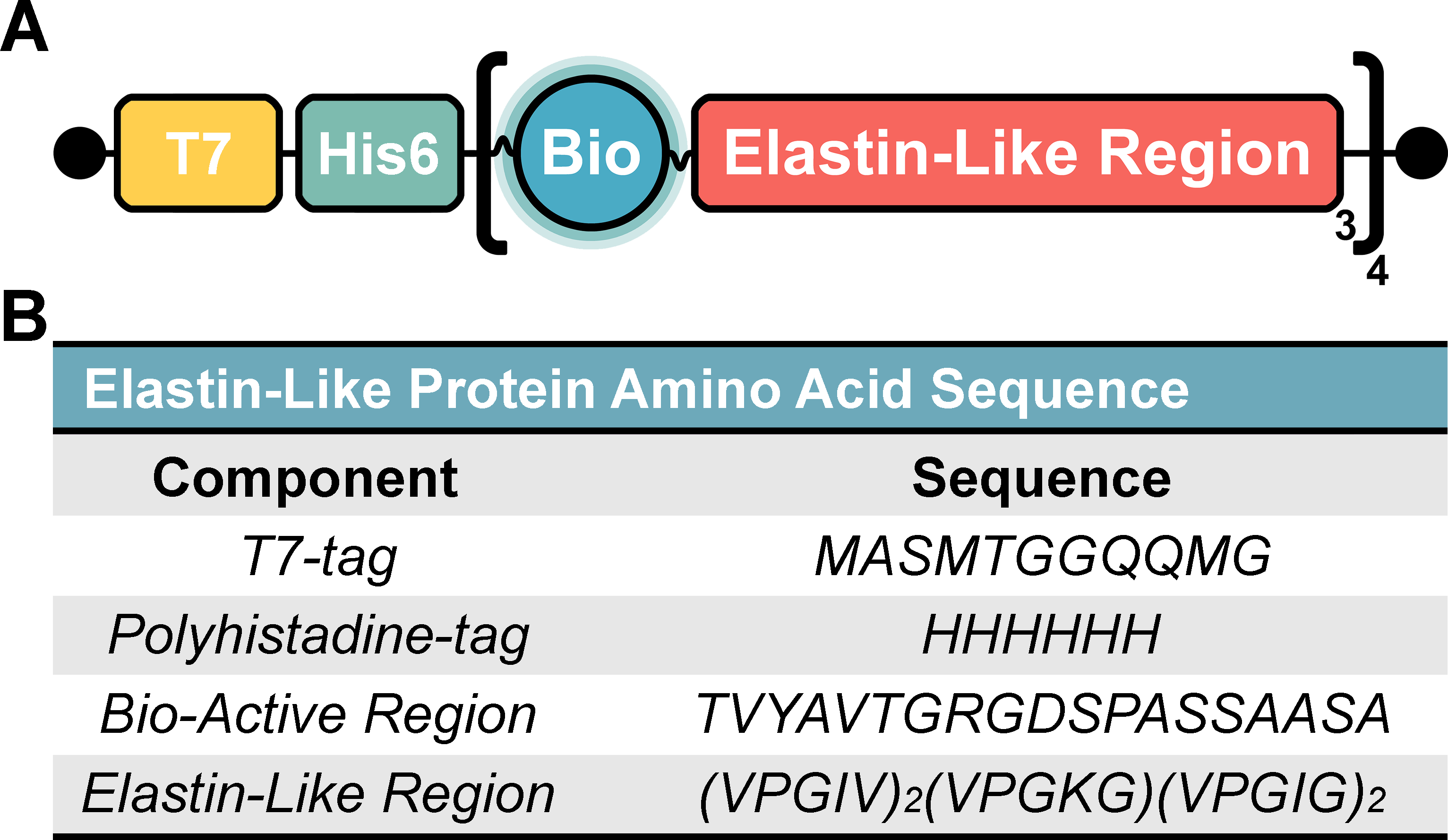

**Figure S1. Elastin-like protein amino acid sequence information:** *(A) A schematic representation of the elastin-like protein (ELP) used in the present manuscript. Our ELP contains three key-components: (1) a tag region with a T7-tag and Polyhistadine-6 tag, (2) a bio-active domain and (3) an elastin-like region. (B) The amino-acid sequence for these components are included for reference.*

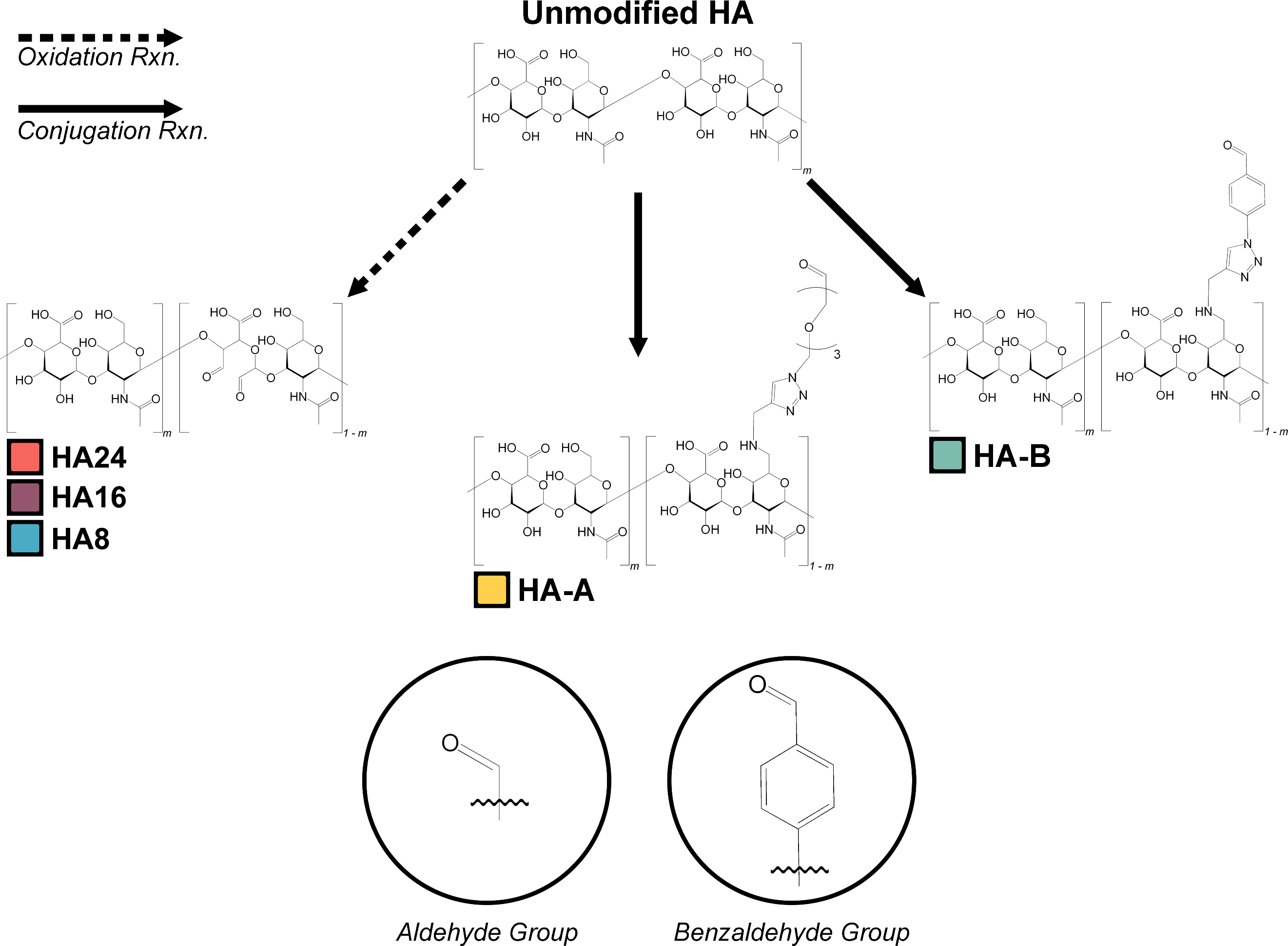

**Figure S2. Summary of conjugation and oxidation reactions for hyaluronan:** *A schematic representation of our different hyaluronan (HA) components. Three variants were produced by oxidizing 1.5-MDa HA for 24 hours (HA24), 16 Hours (HA16), and 8 hours (HA8). The other two HA variants were produced via a 2-part bio-conjugation reaction wherein a 100-kDa HA was first reacted to have 12% of the HA repeat groups modified with an alkyne group (HA-Alkyne). A secondary reaction then conjugated a small molecule with either a pendant aldehyde (HA-A) or benzaldehyde group (HA-B) to the alkyne through a copper-click reaction.*

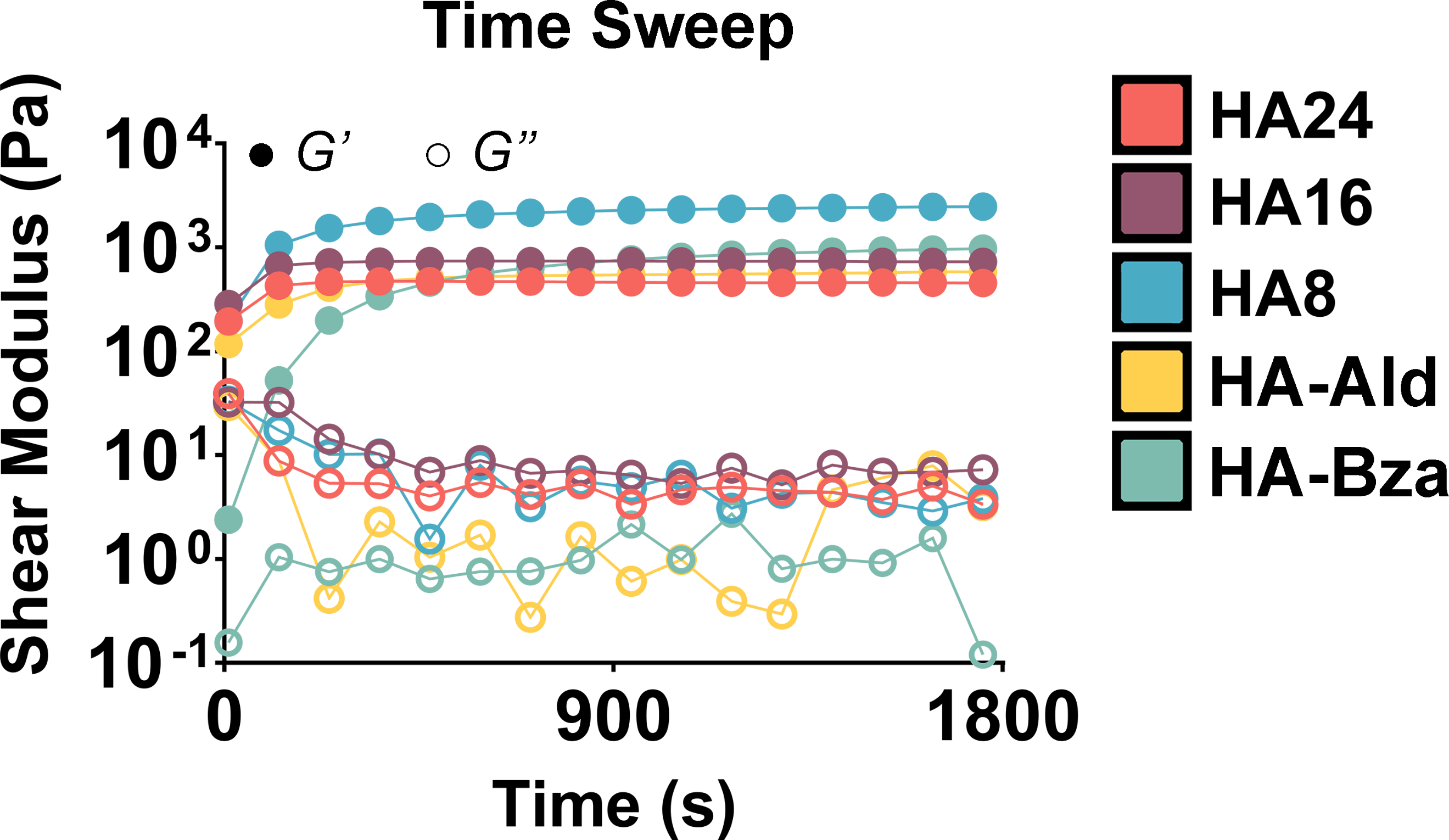

**Figure S3: Representative time sweeps.** *Following mixing, our five gel formulations (all: 2% (w/v) ELP, 1% (w/v) HA) were crosslinked for 30 minutes at room temperature. All five formulations began to gel almost immediately following mixing, and all formulations had a storage modulus (G’) > loss modulus (G’’) prior to the start of measurement. Typical plateau moduli following crosslinking were as follows: ~450 Pa for HA24; ~730 Pa for HA16; ~2500 Pa for HA8; ~590 Pa for HA-A; ~ 980 Pa for HA-B.*

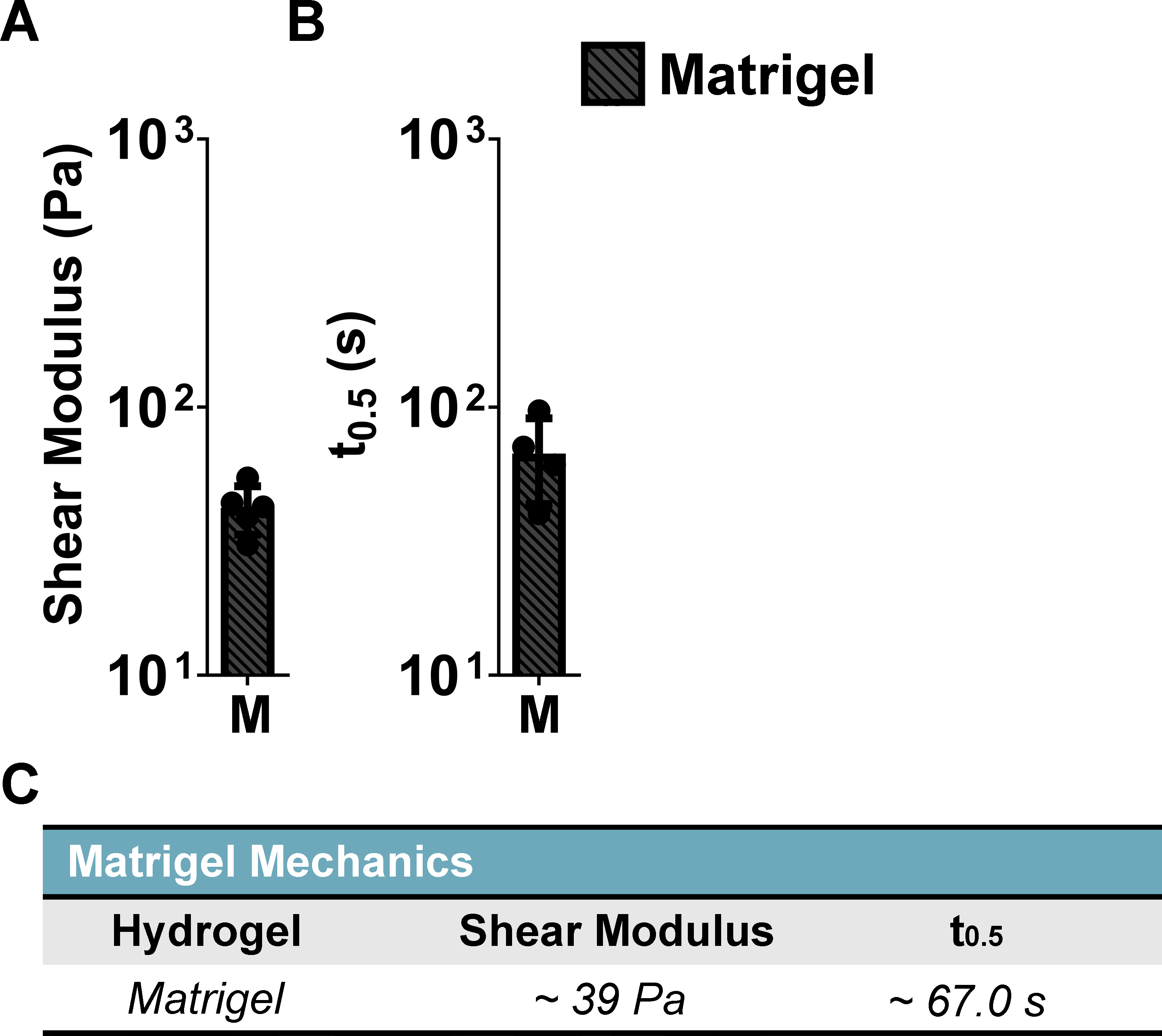

**Figure S4. Summary of animal mechanics:** *Summary mechanics of Matrigel using our prescribed mechanical testing protocols. We found that Matrigel has a shear modulus of ~39 Pa and a stress-relaxation of ~67.0 s.*

***Note: These videos could not be uploaded directly to bioRxiv.org. For inquiries, reach out directly to Riley A. Suhar***

**Figure S5.** **Representative injection videos:** *(A) Representati**ve video of injecting 50 μL of HA24 (dyed red with food coloring for ease of visibility), (B) HA16 (dyed purple with food coloring for ease of visibility), (C) HA8 (dyed blue with food coloring for ease of visibility), (D) HA-A (dyed yellow with food coloring for ease of visibility), and (E) HA-B (dyed green with food coloring for ease of visibility) through a 30-G, manually hooked, needle by hand*

***Note: This video could not be uploaded directly to bioRxiv.org. For inquiries, reach out directly to Riley A. Suhar***

**Figure S6.** **Catheter injection video of HA16:** *To screen the potential for translatability of our hyaluronic acid and elastin-like protein HELP gel system, we tested the injection of 700 µL HA16 (dyed dark green for ease of visibility) through a 150-cm catheter. Injections were done by hand. Note: the beginning of the video has been accelerated (x8). Total injection time is 3 minutes and 8 seconds.*

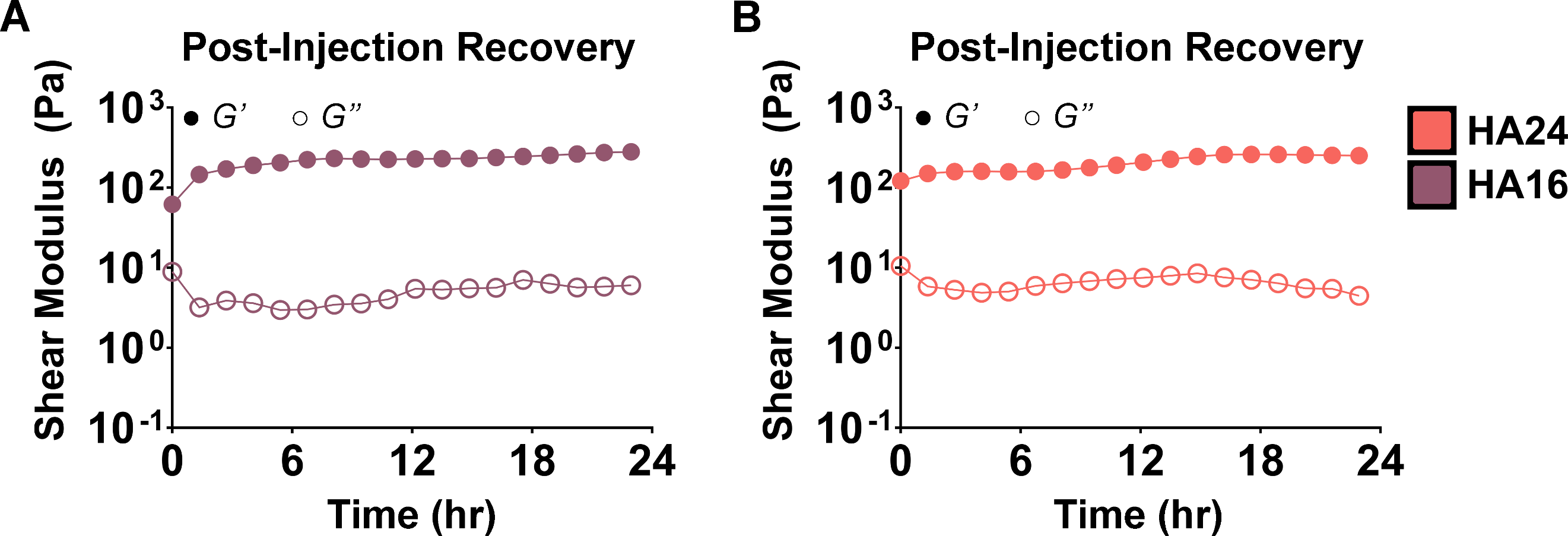

**Figure S7. Recovery experiment for HA16 and HA24:** *24-hour recovery experiment of 50 µL of (A) HA16 and (B) HA24 post-injection at 1% strain and a frequency of 1 rad s^-1^.*

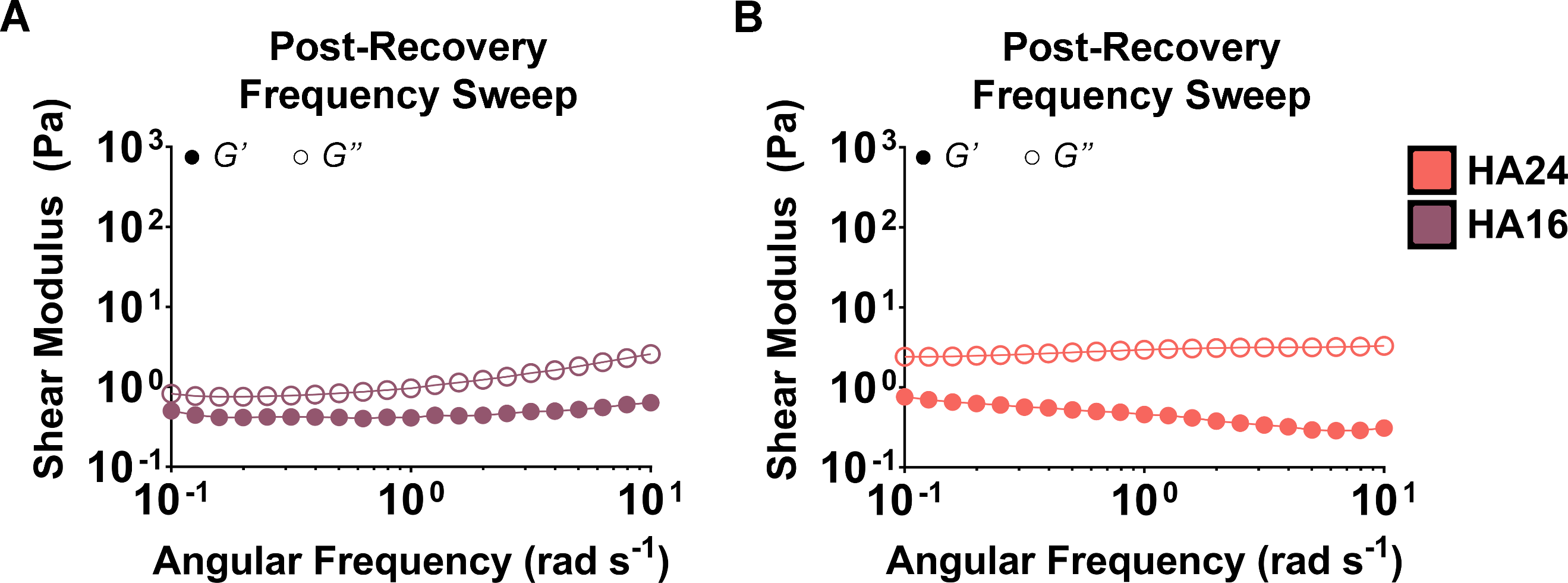

**Figure S8. Post-recovery frequency sweep:** *After 24-hours of recovery post-injection we measured shear stiffness of (A) HA16 and (B) HA24 via a frequency sweep at 10% strain and over a range of 0.1
rad s^-1^ to 10 rad s^-1^, as detailed in the methods section.*

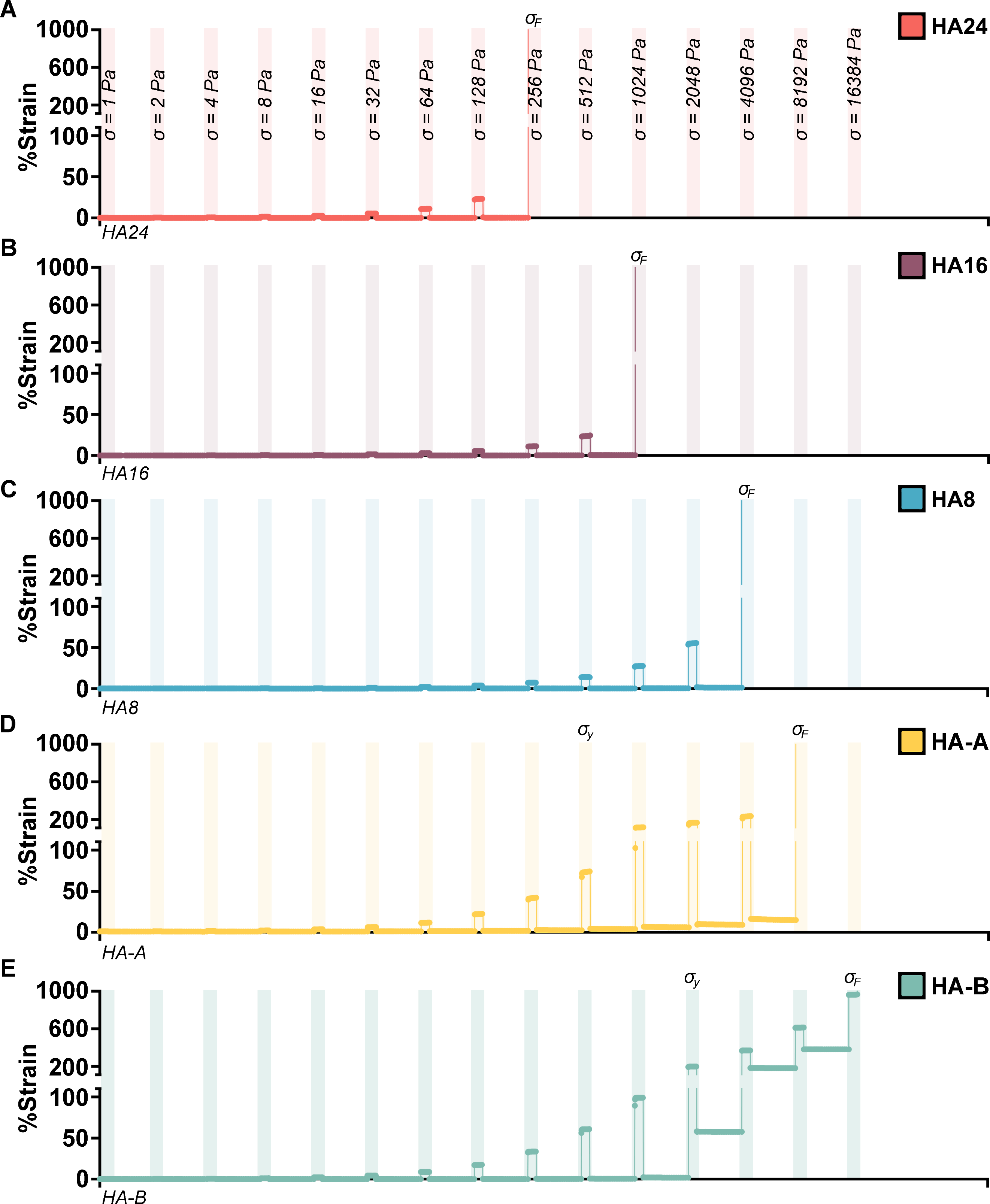

**Figure S9. Representative stress and strain curves:** *Representative failure stress measurement tests for (A) HA24, (B) HA16, (C) HA8, (D) HA-A, and (E) HA-B showing the measured strain over successive stress steps (demarcated by a transparent bar and corresponding stress) and relaxation steps. The stress required to induce fracture is indicated by* $\sigma_{F}$*. Note that the y-axis is broken into two parts: (1) 0 – 100 % strain and (2) 100 – 200% strain. The oxidized HELP (HA24, HA16, and HA8) groups reached much lower strain values (< 50%) prior to failure than the bio-conjugated variants, HA-A and HA-B, which experienced % strain > 100% prior to failure. Additionally, both HA-A and HA-B had evidence of a yielding behavior (% strain >> 0 post-relaxation (*$\sigma= 0$*)) and are denoted with a* $\sigma_{y}$*.*

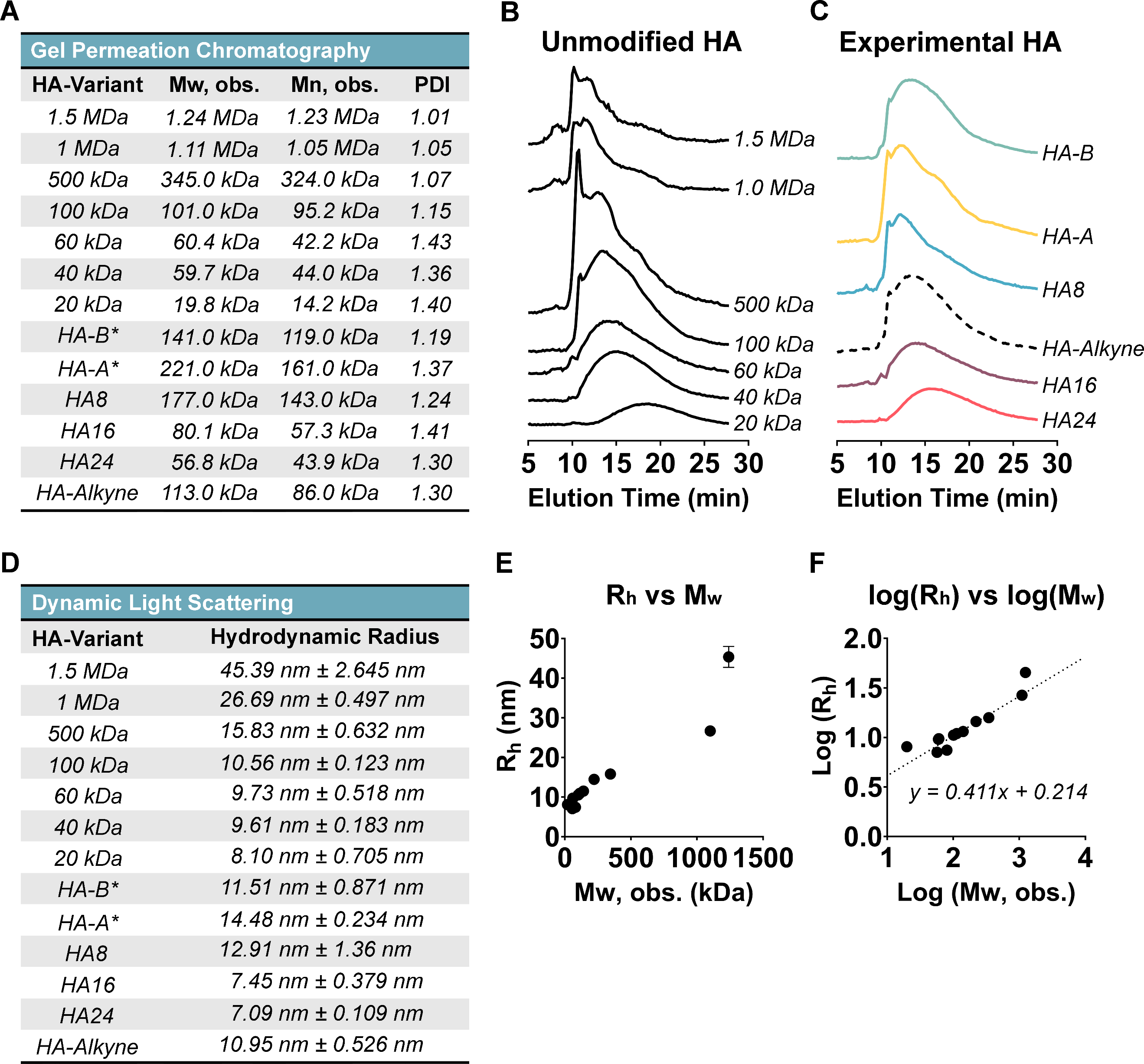

**Figure S10. Estimating molecular weight of modified hyaluronic acid:** *To characterize the molecular weight of our different hyaluronan (HA) variants, we used gel-permeation chromatography (GPC) on both reference, unmodified HA, and our experimental HA groups. (A) Tabulated results showing the observed weight-average molecular weight (Mw, obs.), number-average molecular weight (Mn, obs.), and polydisperisty index (PDI) are shown. GPC measurements were able to accurately determine the molecular weight of our unmodified control groups. HA’s modified by copper click chemistry (HA-B* and HA-A*) appeared to increase in molecular weight, but this is likely due to the presence of bulky side groups increasing their hydrodynamic radii and inflating the measured value. As expected, oxidized HA samples (HA8, HA16, and HA24) decreased greatly from their starting molecular weight (1.5 MDa) as a function of oxidation time. Representative GPC curves have been provided for (B) unmodified HA and our (C) experimental HA groups. Note, we used a specific refractive index increment (dn/dc) value of 0.165 to calculate the approximate values for all samples.* *(D) Using dynamic light scattering (DLS) we measured the hydrodynamic radius of unmodified HA controls and our modified HA groups. The hydrodynamic radius (R_h_) and the Mw, obs. Have been plotted both as (E) linear plots and (F) log-log plots*.

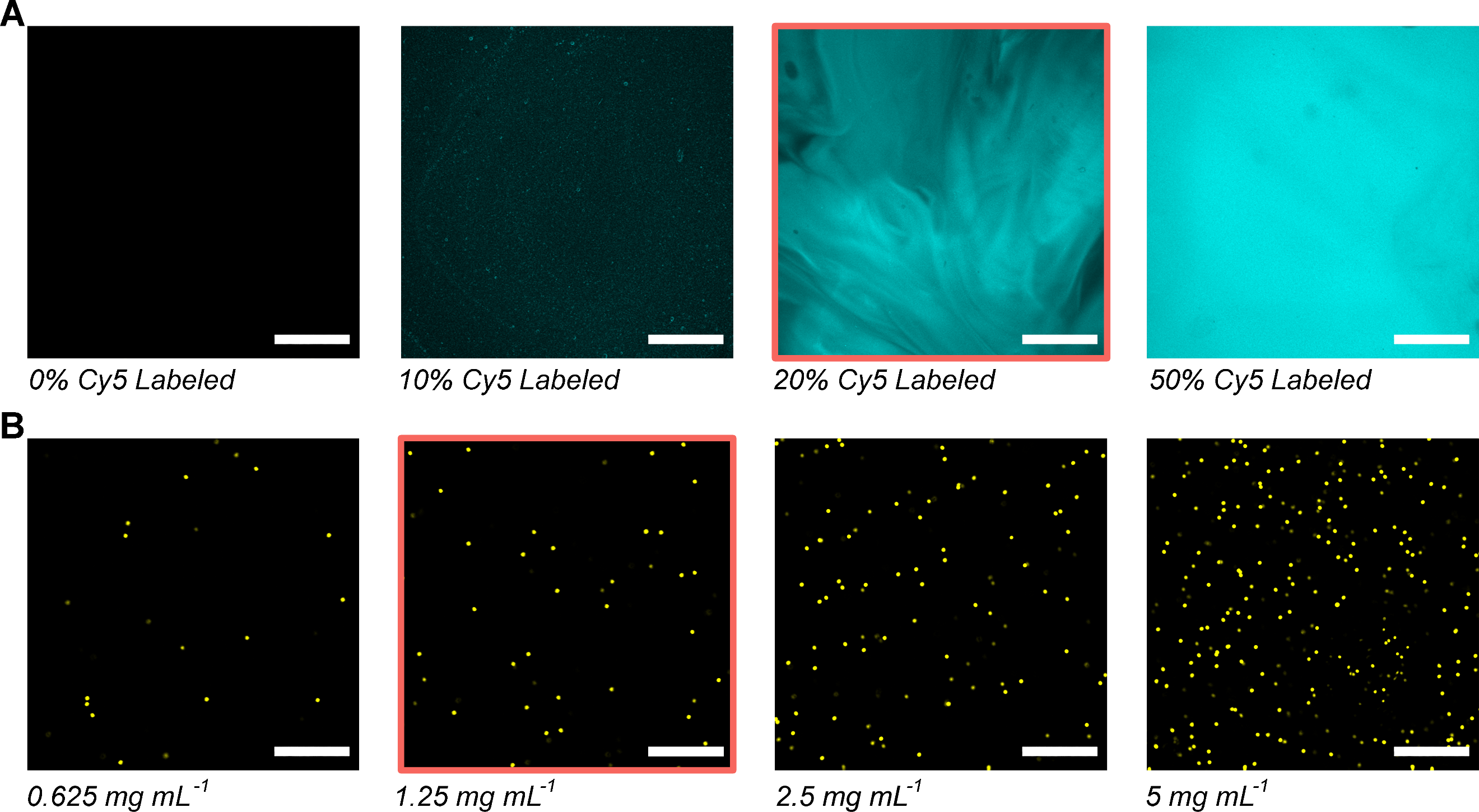

**Figure S11. *Ex vivo* hydrogel fluorescence and microsphere concentration determination:** *To identify an appropriate level of ELP-cyanine-5 labeling and microsphere concentration for subsequent* in vivo *experiments, we prepared HELP gels with varying levels of (A) volume percent labeled ELP and (B) concentration of fluorescent microparticles. From our* ex vivo *screening, we opted to use 20% (v/v) ELP-Cy5-Hydrazine and 1.25 mg mL^-1^, 10-µm fluorescent particles (selections are outlined in red). Based on this image, we further estimated that a 50-µL injection would have approximately ~70,110 microspheres. Note: all images represent 20-µm thick z-stacks to emulate the intended tissue thickness; all scale bars denote 250 µm.*

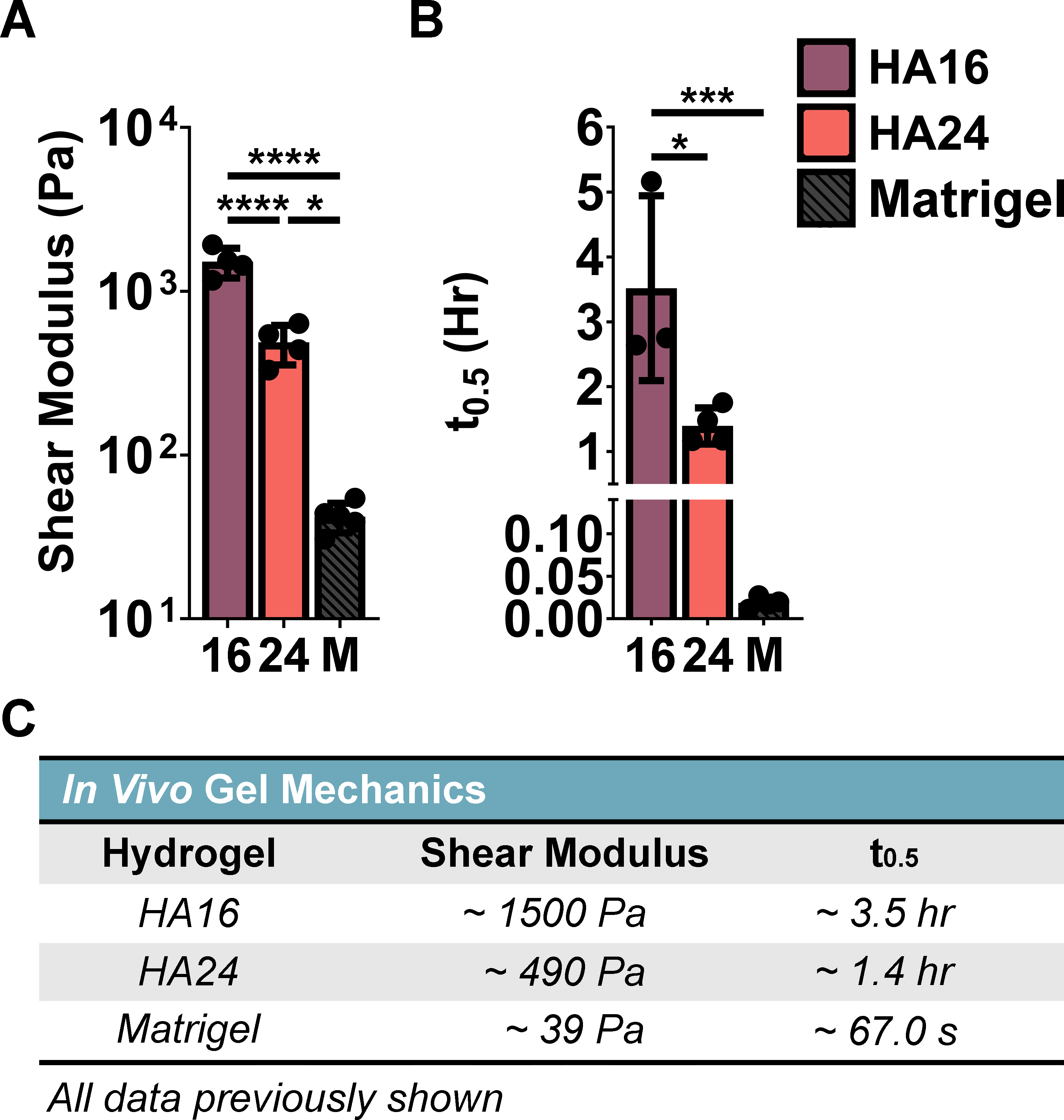

**Figure S12. Summary of animal mechanics:** *Summary of the (A) shear moduli and (B) stress-relaxation of HA16, HA24, and Matrigel. (C) Target values are reported for ease of comparison. Note: this data has previously been shown, but has been recompiled for ease of direct comparison.* *Collectively, these gels established “low” (~40 Pa), “medium” (~500 Pa), and “high” (~1500 Pa) stiffness gels for our* in vivo *retention study. Statistical test: one-way ANOVA, α = 0.05, post-hoc Tukey test. * p < 0.05; *** p < 0.001; **** p < 0.0001.*

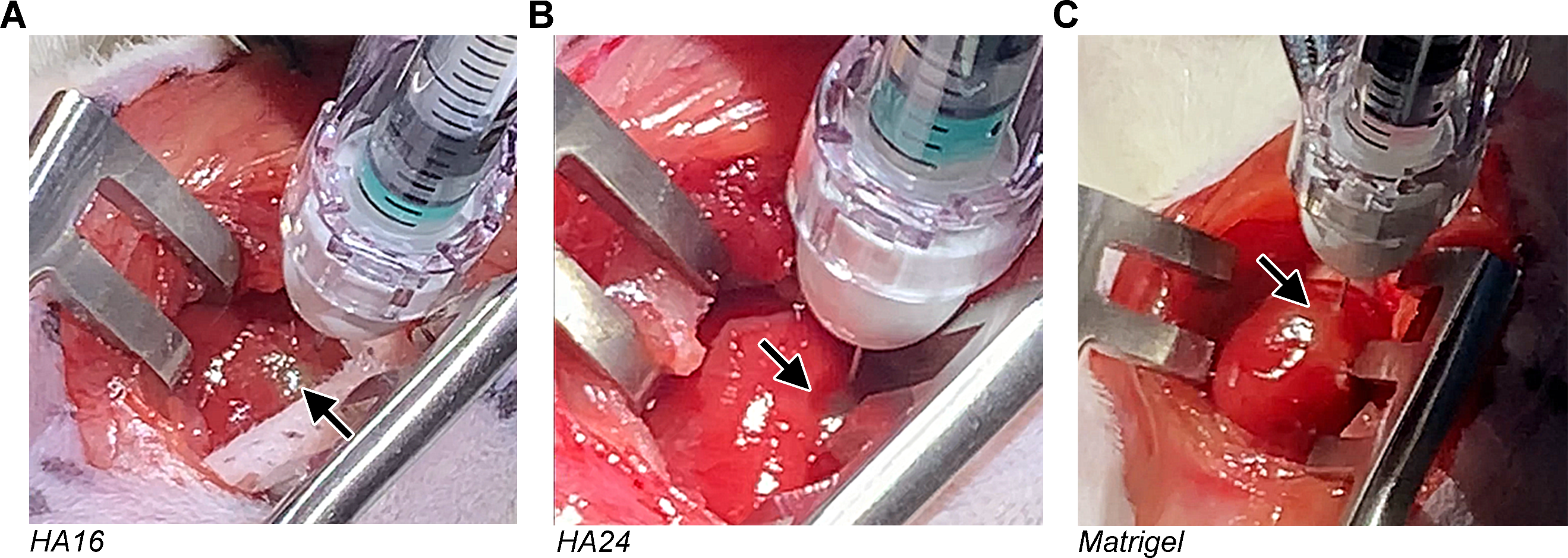

**Figure S13. Representative injection images:** *Representative images showing injections of 50 μL of (A) fluorescently labeled HA16, (B) fluorescently labeled HA24, and (C) Matrigel, each with a final concentration of 1.25 mg mL^-1^ 10-μm fluorescent microspheres. Arrows indicate evidence of material injection of the material, indicated by the formation of a bolus and slight discoloration of the tissue.*

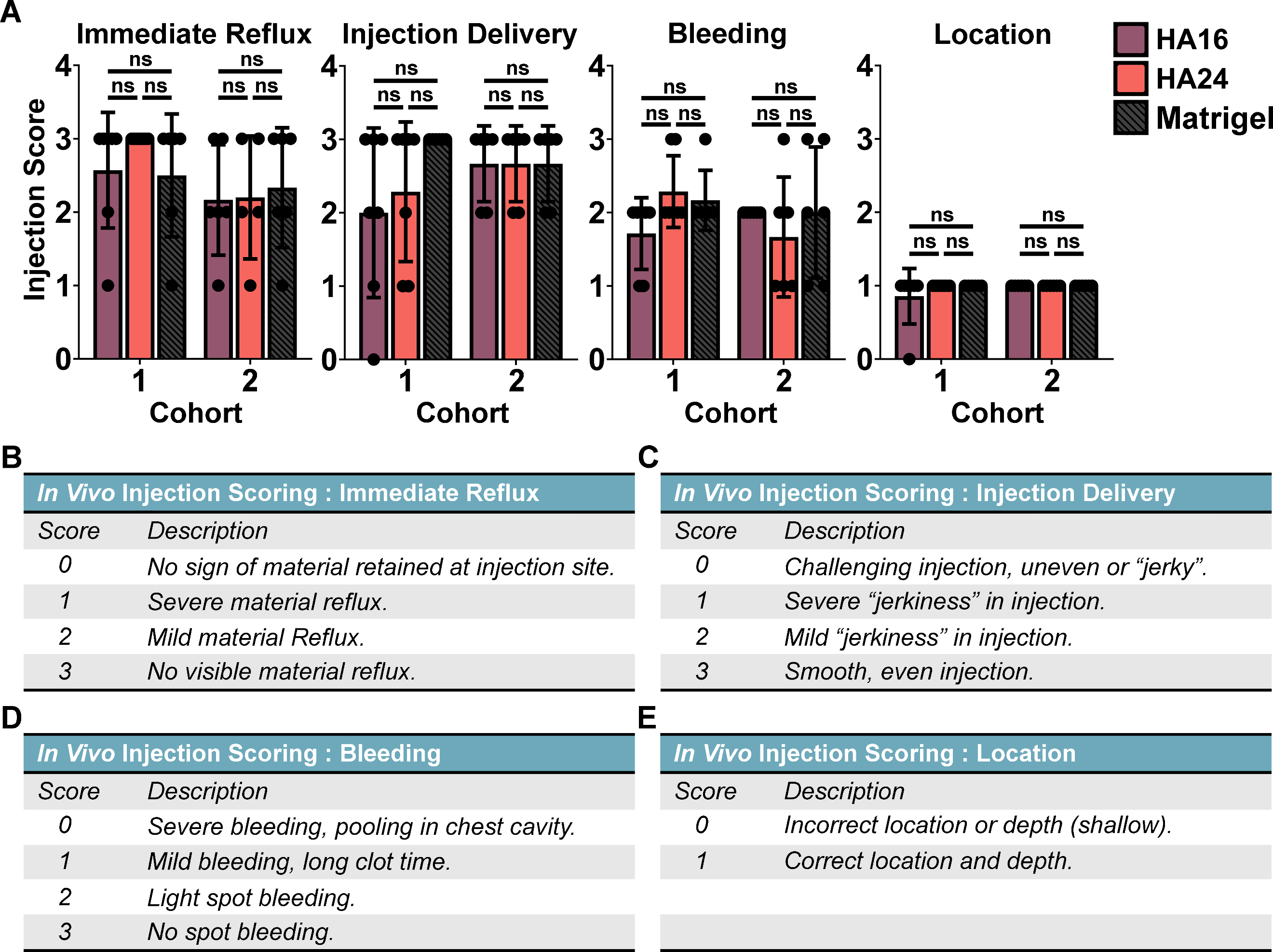

**Figure S14. Injection scoring:** *To probe the possibility that retention could be influenced by the quality of our injections, we devised a 4-part scoring system that looked at: (1) the observable amount of reflux, (2) ease of injection, (3) degree of bleeding post-injection, and (4) correct location of the injection site. (A) These scores were tabulated for each gel group for both cohorts, 1 and 2. Scores were assigned by an external observer, blind to the injection group and time point. (B – D) A breakdown of the scoring scheme has been provided for reference. Notably, we found no significant differences in the quality of our injections between our groups. Statistical test: two-way ANOVA, α = 0.05, post-hoc Tukey test, not significant = n.s.*

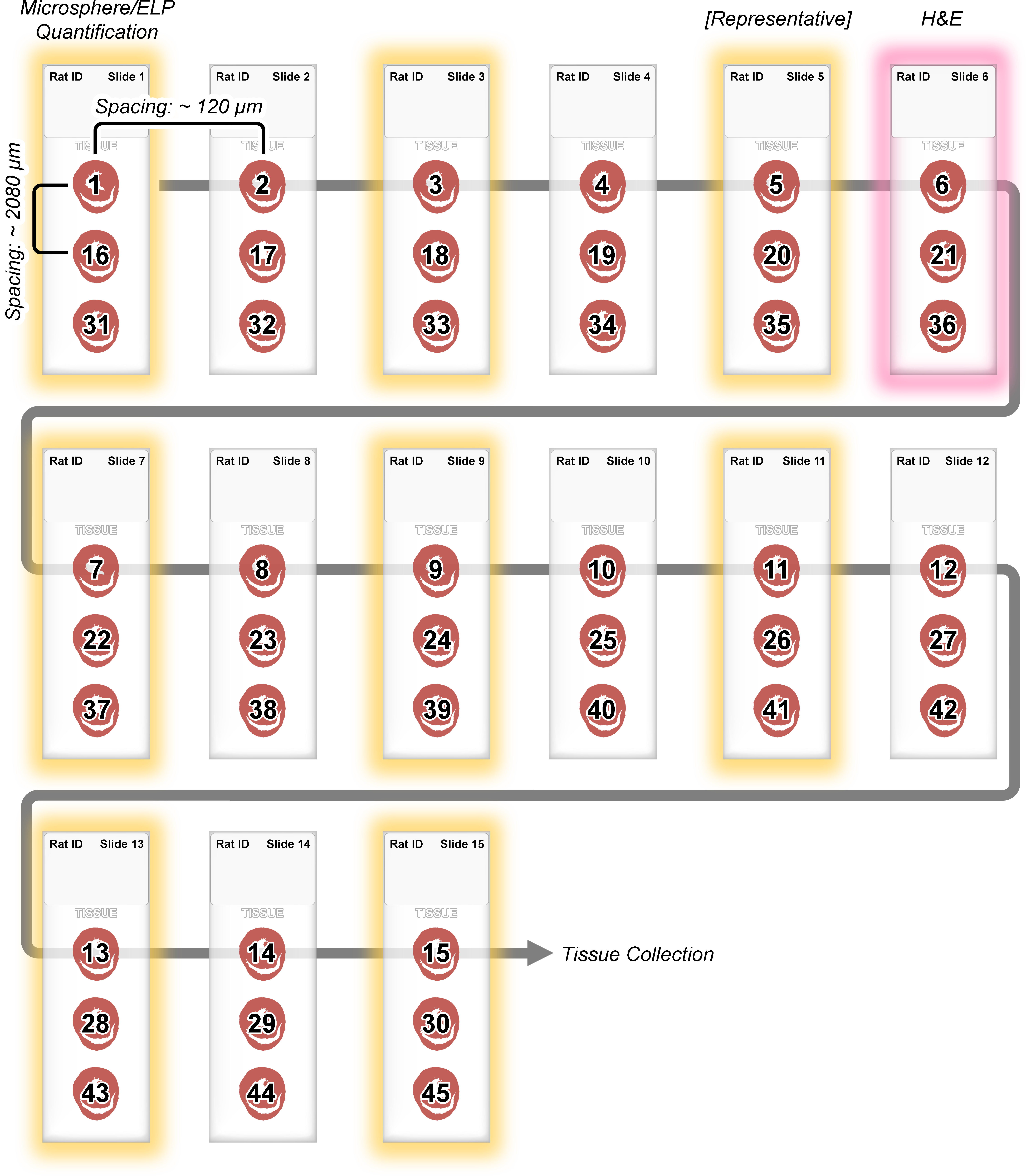

**Figure S15. Tissue sectioning schematic:** *To maintain a consistent and unbiased survey of our different test groups, all tissue was sectioned and sampled according to the above scheme. Briefly, 15 gelatin-coated slides were collected from each animal for analysis. Serial slices (20-μm thick) of heart tissue were made, and every seventh slice was collected; i.e. a slides were prepared from tissue slices spaced approximately ~120-μm apart. Importantly, the exact number of tissue slices between each collected tissue sample was recorded and used in the final quantification. In the above diagram, we have numbered each tissue section (1 – 45) to reflect the general ordering of sample collection, where the first 15 samples (1-15) were placed at the top of each slide, the next 15 samples (16 – 30) were collected in the middle row, and the last 15 were collected on the bottom row (31 – 45). By sectioning tissue this way, each slide spanned approximately 4 mm in height and provided a rapid way of surveying a large amount of tissue with relatively few slides needing to be imaged. From the 15 total slides, 8 (slides 1, 3, 5, 7, 9, 11, 13, and 15) were collected from each animal and imaged for microspheres and elastin-like protein (ELP) according to our described methods. To select a slide for Hematoxylin & Eosin (H&E) staining, we designated from those 8 slides one that was ‘representative’ (having qualities numerically and morphologically indicative of that particular injection) and a slide adjacent to that was then selected for histological staining.*

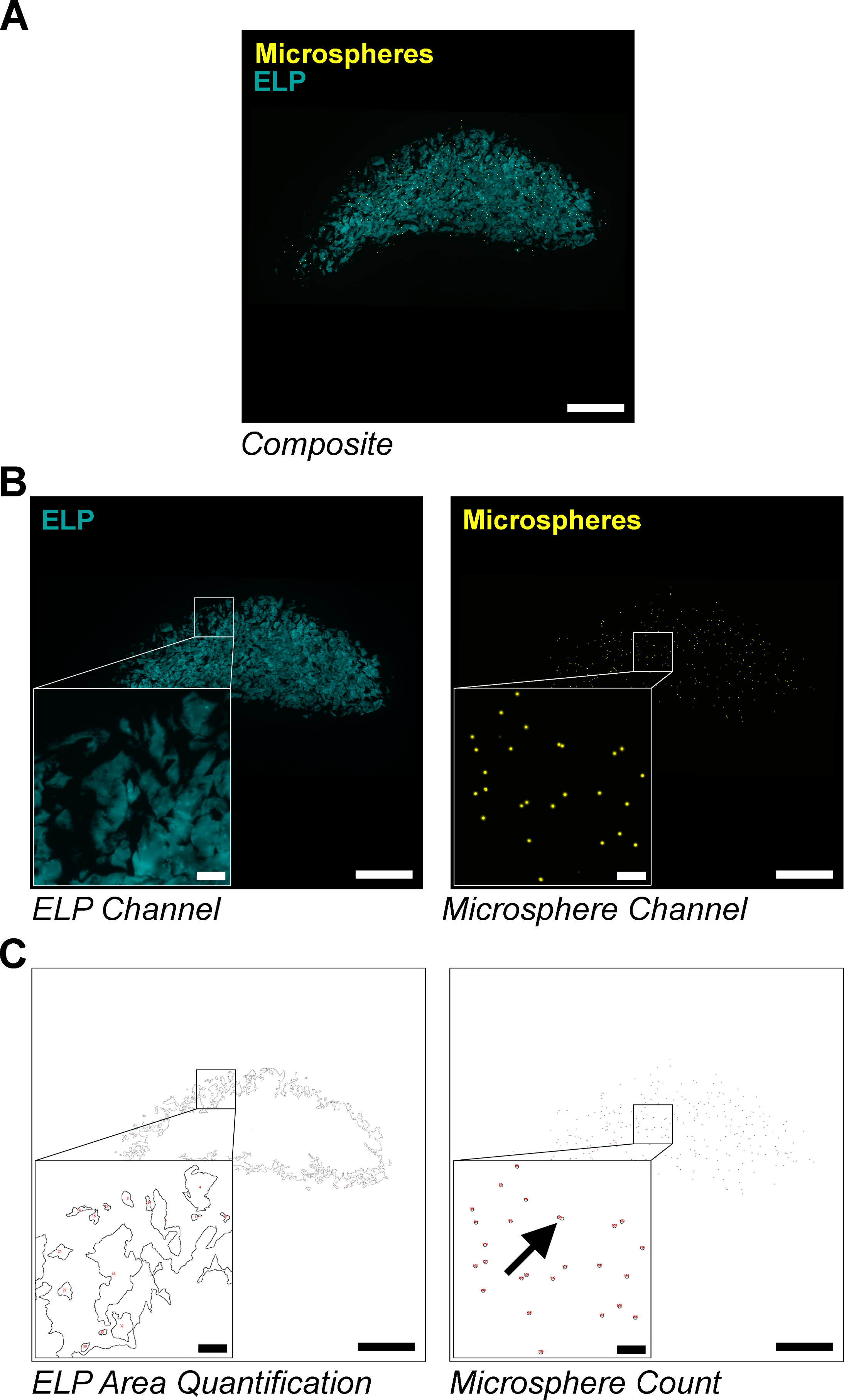

**Figure S16. Example images of microsphere and hydrogel retention quantification:** *Tabulation of elastin-like protein (ELP) cross-sectional area and microsphere count was carried out by automated quantification of fluorescent images. (A) First, 45, 20-µm thick, tissue sections were extracted semi-periodically with a cryostat from a larger 5-mm section of rat heart collected and fixed according to our previously stated methods (see* ***Figure S12****). Tissue sections were imaged with aid of an inverted fluorescent microscope for evidence of injected cargo: ELP and/or fluorescent microspheres. (B) Composite images were then split into either ELP (for HA16 and HA24) or microsphere channels (HA16, HA24, and Matrigel) and saved separately. (C) These images were then converted to binary images (using a constant threshold value) and either the area of ELP or the number of microspheres were tallied for each slice using a particle counter in ImageJ. To generate more accurate counts, tallies from each image slice were then checked for instances of ‘clustering’ (multiple microspheres being counted as one; black arrow) by a secondary Python script that divided the cross-sectional area of each microsphere by the average area of an individual microsphere, and extra counts were added accordingly for each slice. Note: scale bar on low magnification images represents 1 mm, scale bar on high-magnification images represents 100 µm.*

***Note: This file could not be uploaded directly to bioRxiv.org. For inquiries, reach out directly to Riley A. Suhar (******)***

**Figure S17. Python script for post-processing microsphere count:**  *A Python script was written and used to account for microsphere ‘clustering’ which superficially undercounted the number of microspheres in our captured images.*

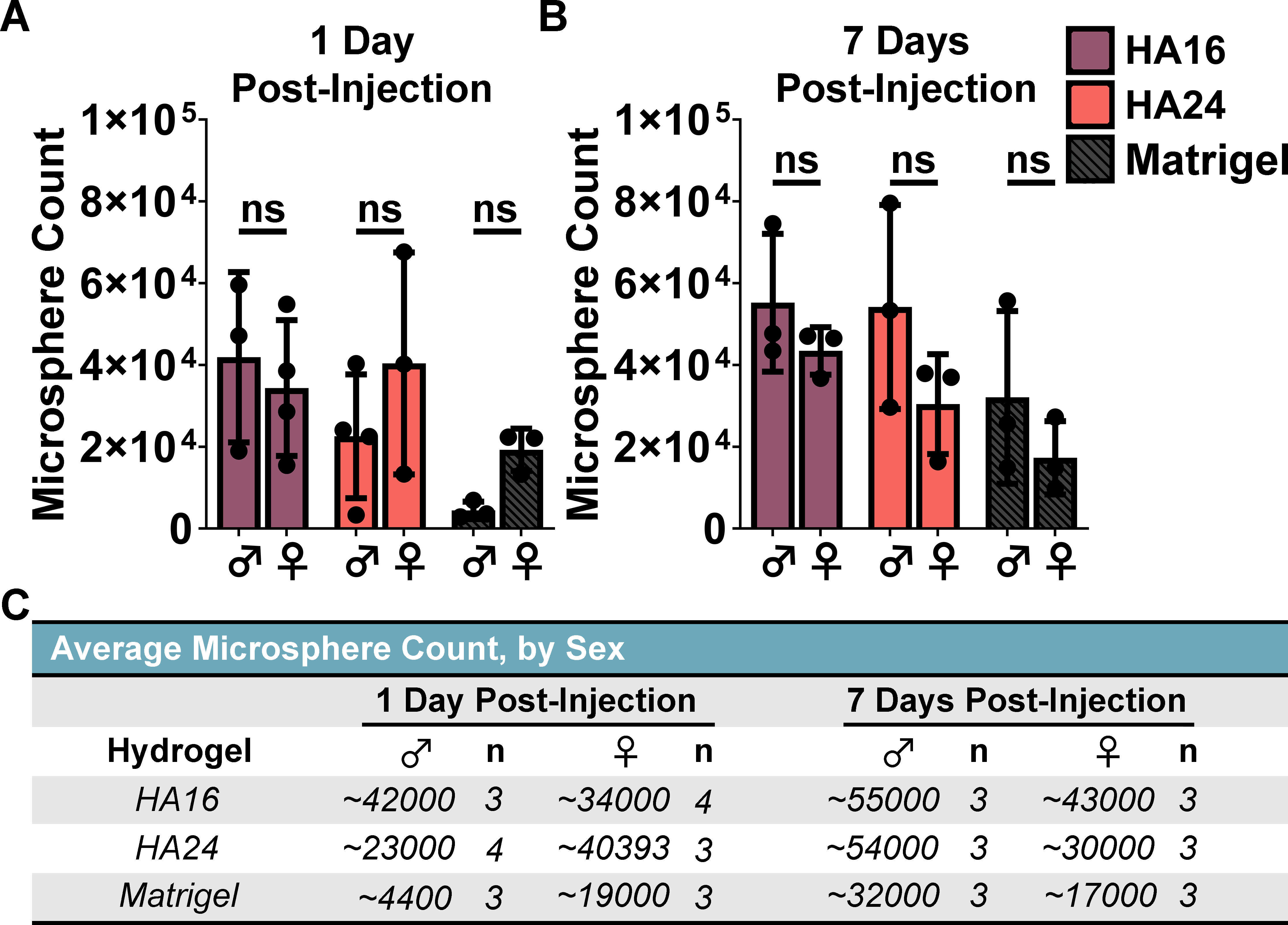

**Figure S18. Sex-based differences in microsphere retention:** *To investigate the possibility of any sex-related differences, we plotted the average microsphere count for each hydrogel, by sex, for both (A) 1 Day and (B) 7 Days post-injection. Importantly, we did not observe any significant differences based on sex. Generally, it appeared that male rats had a greater degree of retention, but not in a statistically significant manner. (C) The results have been additionally summarized for reference.* *Statistical test: two-way ANOVA, α = 0.05, post-hoc Tukey test, not significant = n.s.*
